# Upregulation of the Unfolded and Mitochondrial Unfolded Protein Responses in Oxidative Stress-Induced Cataract

**DOI:** 10.64898/2026.08.19.745850

**Authors:** Sarah R. Zelle, W. Hayes McDonald, Hassane S. Mchaourab, Kevin L. Schey

## Abstract

**Purpose:** Oxidative stress is thought to contribute to the development of age-related cataracts (ARCs), but the mechanisms by which oxidative damage leads to the opacification of the lens remain unclear. Previous studies suggest that oxidative stress can disrupt lens proteostasis. Therefore, it was hypothesized that ARCs arise from proteomic changes driven by an age-associated decline in oxidative stress defenses that interact with the lens proteostatic state. To test this hypothesis, proteomic analyses of lenses exposed to oxidative stress were performed to examine oxidative and proteostatic stress responses *in vivo*.

**Methods:** Cataract formation was induced by injecting hydrogen peroxide into the aqueous humor of adult zebrafish. *nrf2^fh318/fh318^* zebrafish were used to model the reduced oxidative stress protection observed in aged human lenses, while *cryaba^-/-^* zebrafish were used to model impaired lens proteostasis. Resulting opacities in WT, *cryaba^-/-^*, *nrf2^fh318/fh318^*, and *cryaba^-/-^; nrf2^fh318/fh318^* lenses were quantified and proteomic changes in the cortex were analyzed using data independent acquisition Parallel Accumulation Serial Fragmentation mass spectrometry.

**Results:** Hydrogen peroxide treatment induced the formation of cortical cataracts. Proteomic results showed that, dependent on genotype and day, oxidative stress activates the unfolded and mitochondrial unfolded protein responses. Additional changes were also observed in energy metabolism, Ca^2+^ homeostasis, protein degradation, and cytoskeletal and extracellular matrix remodeling pathways.

**Conclusions:** Treated zebrafish lenses successfully model ARC and mass spectrometry proteomics identified the unfolded and mitochondrial unfolded protein responses as potential therapeutic targets for ARC.

## Introduction

Cataracts, where the ocular lens becomes opacified and scatters light, are the primary cause of blindness globally.^1–4^ The most common type of cataracts are age-related cataract (ARCs).^5^ Oxidative stress is considered a major contributor to ARC formation, and reactive oxygen species, such as hydrogen peroxide (H_2_O_2_) have been shown to disrupt the lens proteome.^6,7^ Nonetheless, a thorough understanding of the molecular determinants of the lens oxidative stress response and how it results in ARC formation has yet to be determined.

Nuclear factor erythroid 2-related factor 2 (Nrf2) is an active lens transcription factor considered the master regulator of the oxidative stress response, and Nrf2 deficient-models have previously been used to study ARCs.^8–10^ For instance, aged *Nrf2^-/-^* mice can develop cataracts in all major regions of the lens.^11^ Outside of the lens, Nrf2 has also been characterized using models such as *nrf2^fh318/fh318^* zebrafish, which carry a point mutation that impairs Nrf2 DNA-binding activity, reducing expression of Nrf2-controlled transcripts. As a result, larvae are sensitive to treatment with oxidative stress inducers.^12^ Under oxidative stress conditions, Nrf2 translocates to the nucleus to bind antioxidant response elements and upregulates transcription of antioxidant genes.^8–10^ In addition to lens antioxidants, there are also highly abundant small heat shock proteins, such as α-crystallins, that bind partially denatured proteins to prevent aggregation, and act as proteostatic sensors.^13,14^ Mishra et al. utilized CRISPR/Cas9 to create a loss of Cryaba function zebrafish line, *cryaba^-/-^*, and discovered that Cryaba is involved in stress resistance in the heart and lens.^15^

To investigate the role of the oxidative stress response in the formation of ARCs, Park et al. utilized *nrf2^fh318/fh318^* zebrafish as a model for ARC, mimicking the reduced oxidative stress protection seen in aged lenses.^16^ Furthermore, to examine how the oxidative stress response interacts with lens proteostatic pathways, the authors also employed *cryaba^-/-^* zebrafish as a mimic for perturbed lens proteostatic state. Park et al. observed significantly more lens defects in *cryaba^-/-^* embryos compared to WT, whereas *cryaba^-/-^; nrf2^fh3^*^18^*^/fh318^*embryos were comparable to WT and *nrf2^fh318/fh318^* embryos. This finding suggests that the oxidative stress response alters the lens proteostatic state to reduce the number of zebrafish embryos with lens defects. However, the proteomic changes underlying these results were not examined, nor were the lenses of adult zebrafish exposed to oxidative stress.

To evaluate the *nrf2^fh318/fh318^* and *cryaba^-/-^* genotypes in the context of oxidative stress, the eyes of adult zebrafish were injected with phosphate buffer saline (PBS) as a control or H_2_O_2_ to induce cataract formation. Prior et al. first reported that injection of H_2_O_2_ into the eyes of adult WT zebrafish induced cataracts 1 day post-H_2_O_2_ treatment, which resolve over the course of 22 days post-injection.^17–20^ In the current study, the data-independent acquisition parallel accumulation-serial fragmentation (diaPASEF) mass spectrometry proteomics methodology described in Zelle et al. was employed to examine lens proteome changes after H_2_O_2_-inducedcataract formation in adult zebrafish.^21^

## Materials and methods Materials

All reagents and materials were purchased from Sigma-Aldrich (St. Louis, MO) and Fisher Scientific (Waltham, MA) unless indicated otherwise. S-Trap micro columns were procured directly from ProtiFi (Farmingdale, NY).

### Zebrafish maintenance

AB-WT (WT), *cryaba^-/-^*, *nrf2^fh318/fh318^*, and *cryaba^-/-^; nrf2^fh318/fh318^* zebrafish (*Danio rerio*) embryos were raised to 3-9 months of age at the Vanderbilt Zebrafish Aquatic Facility. Zebrafish were kept on a 14:10 hour light/dark cycle and in deionized water with 30 mg/L instant ocean at 28.5 °C. All experiments involving zebrafish were authorized by the Vanderbilt University Institutional Animal Care and Use Committee and adhere to the ARVO Statement for the Use of Animals in Ophthalmic and Vision Research.

### PBS and H_2_O_2_ injections into the zebrafish eye

Injections to induce the formation of a cataract in adult zebrafish were carried out following previously described methods.^17–20^ A Sutter Instrument P-97 micropipette puller with settings 300 PSI, 315 heat, 70 pull, velocity 65, and 80 msec was used to pull 3.5 in long Drummond capillaries with a 1.14 mm outer diameter. Zebrafish were anesthetized using Tricaine Methanesulfonate and wrapped in a damp paper towel. A sterile 30G insulin needle was used to puncture the cornea and a Drummond Nanoject II Automatic Nanoliter Injector outfitted with a pulled glass capillary taped to a zebrafish microinjector apparatus was used to administer 0.483 µL of either 1X PBS or 2.5% H_2_O_2_ in 1X PBS into the aqueous humor via the corneal puncture.

### Dissection and imaging

An Eschenbach Mobilux LED hand-held magnifier (12.5X magnification) and an iPhone 13 Pro at 2X magnification with the default camera settings was used to take images of the intact eyes 3 and 7 days post-injection.^22^ Fourteen lenses for each genotype (4 and 8 days post-PBS or post- H_2_O_2_ injection), except for the *cryaba^-/-^* 4 days post-PBS treated group, which had thirteen lenses, were dissected and placed in a 35 mm glass-bottom cell culture dish in PBS.^16^ A power analysis using G*Power 3.1.9.7^23,24^ (effect size=0.25, power=90%, error probability=0.05) indicated a minimum sample size of 171, or ∼11 samples per condition was needed for treatment and day comparisons (numerator degrees of freedom=1). A minimum sample size of 231, or ∼14 samples per condition was needed for genotype comparisons (numerator degrees of freedom=3). Images were taken in darkfield using a Zeiss Axio Zoom V16 microscope at 40X magnification with 35 msec exposure time.

### Analysis of images

Lens images were processed using custom Fiji macros in **Supplemental Files 1 and 2**. Each raw image was converted to 8-bit and underwent local contrast and median filtering. PBS and H_2_O_2_ images were then thresholded separately to generate binary masks of the lens. Particle analysis was applied to distinguish the lens from the background and a distance map was used to determine the maximum radius of each lens. The mean gray value was measured in concentric 10 µm-thick rings extending from the center to the edge of the lens. R-scripts were used to normalize the radial distance to account for slight differences in lens size. For visualization, replicates were fitted using a natural cubic spline and averaged together. The area under each replicate’s spline was calculated and a three-way Type III ANOVA was performed using log-transformed values. Post-hoc estimated marginal means (emmeans) were compared using Tukey-adjusted pairwise contrasts for genotype and nominal contrasts for treatment and day using the car and emmeans packages in R.

### Preparation of samples for discovery mass spectrometry proteomics analysis

The zebrafish lens nucleus and cortex were separated as in Zelle et al.^21^ The lens was placed into the neck of a Biotik 20 µL xTIP4 pipette tip in a droplet of 100 mM Tris pH 7.8, 1.5 mM ethylenediaminetetraacetic acid (EDTA) buffer and spun at 5,000 g to shear off the cortex region. The intact nucleus in the pipette tip was then freed using a razor blade, washed, and both regions were snap-frozen. Notably, not all H_2_O_2_-treated lenses could be successfully separated. Therefore, only lenses that were separated successfully were retained for further analysis.

Of the fourteen, or thirteen for *cryaba^-/-^* at 4 days post-PBS treatment, intact lenses imaged and separated for each sample type, four cortex replicates were thawed and prepared as in Zelle et al.^21^ 25 µL of 100 mM Tris pH 7.8, 1.5 mM EDTA, 10% sodium dodecyl sulfate (SDS) buffer was added to each sample. Samples were homogenized using a handheld electric pestle and spun down. Protein concentration in the supernatant was measured using a bicinchoninic assay before being reduced, alkylated, and digested on S-Trap micros.

### Data acquisition and analysis

Peptides were resuspended in 0.2% FA and adjusted to 50 ng/μL with 0.015% n-dodecyl-ß-D- maltoside. diaPASEF data were acquired using a custom acquisition method on 2 μL of block randomized samples following the LC-MS conditions established in Zelle et al.^21^ Briefly, peptides were resolved via a 30-minute aqueous to organic gradient, approximately 45-minute total cycle time, on a 25 cm x 75 µm PepSep column coupled to a Bruker timsTOF HT instrument.

Protein identification was performed with DIA-NN 2.0^25,26^ using the default library-free settings as in Zelle et al.^21^ A UniProt Swiss-Prot and TrEMBL zebrafish database (UP000000437, downloaded 02/10/2025, 26,712 entries) was utilized and MS1 and MS2 mass tolerances were fixed to 15 ppm. R scripts were used to analyze data for peptides that had a q-value < 1% and corresponded to at least two unique peptides per protein group. For each peptide, the median intensity across all runs was calculated and the fold change relative to the median was determined. Run-specific median fold changes were then calculated using only peptides detected in at least 80% of all runs and individual peptide intensities were divided by the corresponding run-specific median fold change (**Supplemental Figure 1**).

Protein abundances were calculated from normalized precursor intensities using the diann_maxlfq function from the diann package (**Supplemental File 3**).^25^ Values were then log_2_- transformed and statistical testing was performed using limma.^27^ Only proteins detected in at least three of four replicates per sample type were included. Due to the high biological variability in zebrafish, un-adjusted p-values were used for exploratory analysis, and results were plotted using the EnhancedVolcano package.^28,29^ Proteins absent in all replicates of a group but present in at least three out of four replicates of the other group and had an average mean intensity greater than or equal to the overall median intensity of all identified proteins were excluded from significance testing and retained in a separate list (**Supplemental File 4**). Gene Set Enrichment Analysis (GSEA) was completed using the WebGestaltR package and the enrichment significance was determined using a false discovery rate (FDR) cut-off of < 0.05.^30,31^ All figures were generated using the ggplot2 package. Only proteomic data from 4 days post-treatment is discussed in this article, but all data and R scripts used for analysis are available via ProteomeXchange with identifier PXD074656.

## Results

### H_2_O_2_-injections induce cataract formation

H_2_O_2_ injections into the aqueous humor of the adult zebrafish resulted in the formation of a cortical cataract in all groups as seen via an opaque ring around the center of the lens 4 days post-injection (**Figure 1A and 1B**). H_2_O_2_ injected eyes also expanded from the socket immediately after injection, normalizing by 1 day post-injection (**Supplemental Figure 2**). Notably, only H_2_O_2_- injected eyes exhibiting this expansion developed cataracts.

**Figure 1.**
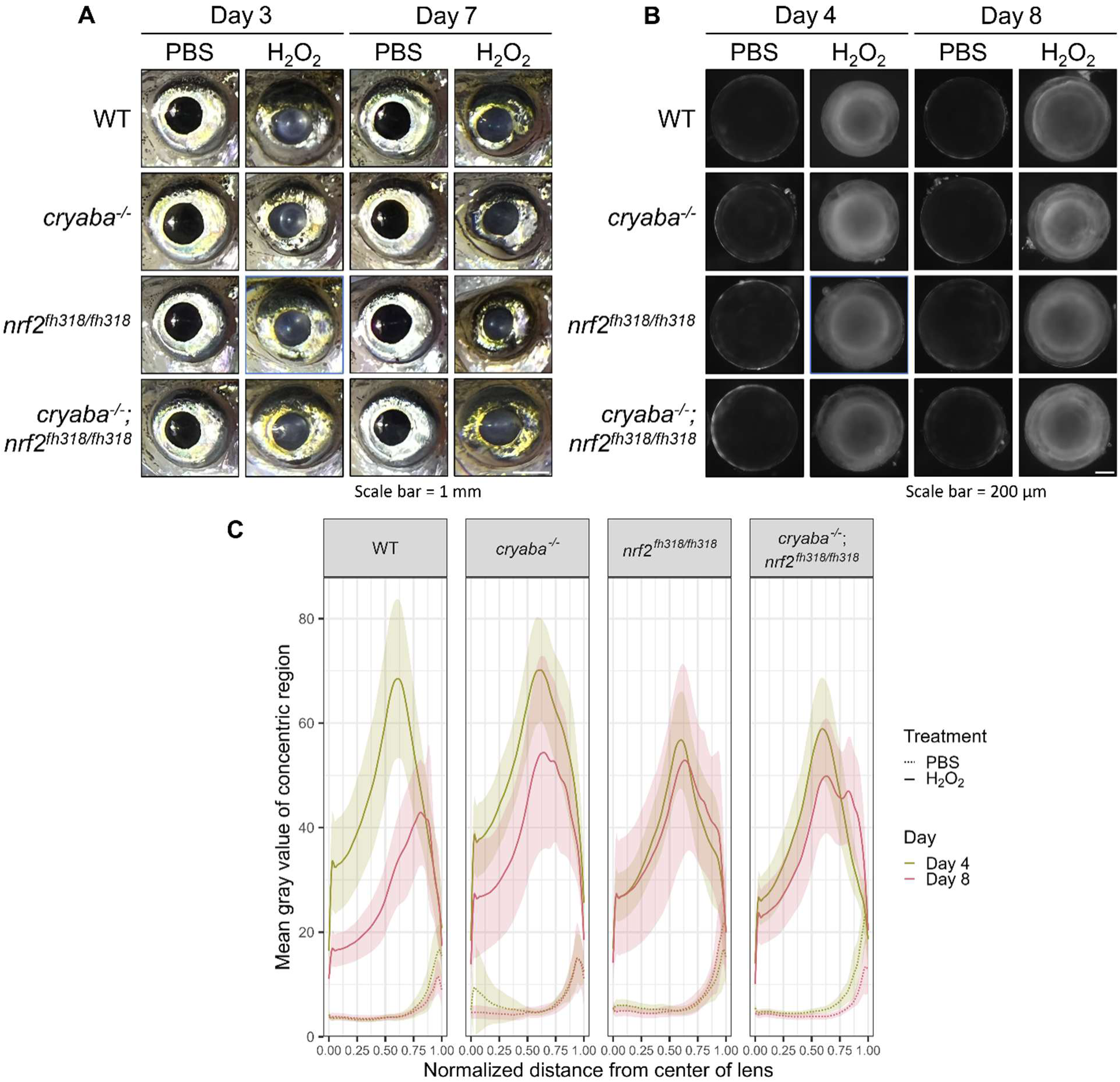
Representative images of A) intact eyes 3 and 7 days post PBS or H_2_O_2_ injection and B) dissected lenses 4 and 8 days post PBS or H_2_O_2_ injection. C) Mean gray values from the normalized center of the lens for WT, *cryaba^-/-^*, *nrf2^fh318/fh318^*, and *cryaba^-/-^*; *nrf2^fh318/fh318^* lenses 4 and 8 days post PBS or H_2_O_2_-treatment. All PBS injected eyes remained clear and all H_2_O_2_ injected eyes were opaque. The white scale bar for the intact eyes corresponds to 1 mm. All PBS injected lenses remained clear and all H_2_O_2_ injected lenses exhibited a ring-shaped cataract. The white scale bar for the dissected lenses corresponds to 200 µm. Replicates were fitted to a natural cubic spline and averaged together and visualized with 95% confidence intervals. The H_2_O_2_-treated lenses exhibit an increase, followed by a decrease in opacity from the normalized distance from the center of the lens. The PBS-treated lenses had an overall lower mean gray value and thus lower opacity compared to the H_2_O_2_-treated lenses. The increase in opacity at the extremities of the lens in the PBS-treated are likely due to artifacts from the edge of the lens.

The opacities in the treated lenses were quantified using mean gray values measured at a distance relative to the center of the lens (**Figure 1C**). Statistical testing revealed significant main effects of treatment, genotype, and day on lens opacity, but no significant interaction terms (**Supplemental Table 1**). Optional post-hoc tests confirmed that H_2_O_2_ treatment significantly increased lens opacity compared to PBS treatment (**Supplemental Table 2**), and 4 days post- treatment lenses were significantly more opaque than 8 days post-treatment, for all conditions pooled together (**Supplemental Table 3**). A Tukey-adjusted post-hoc test revealed that WT lenses had a significantly lower opacity than *cryaba^-/-^* and *nrf2^fh318/fh318^* lenses when pooling across all treatments and days, and the *cryaba^-/-^*; *nrf2^fh318/fh318^*lenses did not differ significantly from WT lenses (**Supplemental Table 4**).

### Proteomic analysis of WT vs. mutant lenses 4 days post-PBS treatment

Analysis of the zebrafish lens cortex identified an average of 40,480 peptides and 3,516 protein groups per sample type after filtering. H_2_O_2_-treated lenses had a an average of 45,166 peptides and 3,906 protein groups per sample, whereas PBS-treated controls averaged 35,792 peptides and 3,126 protein groups per sample.

Compared to WT lenses, *cryaba^-/-^* lenses exhibited a decreased abundance of proteins involved in the lens microcirculation system, cell adhesion, and cytoskeleton, together with an increased abundance in extracellular matrix (ECM) proteins. Results from GSEA showed GO terms associated with metabolism, catabolism, and translation were significantly enriched (**Supplemental Figure 3** and **Supplemental Tables 5 and 12**). Compared to WT, *nrf2^fh318/fh318^*lenses showed a significant decrease in Calb1, a cytosolic Ca^2+^ buffering protein, Cst14b.2, a cysteine protease inhibitor, and cytoskeletal components. Notably, Atp2a3, a Ca^2+^-ATPase that pumps calcium from the cytosol into the endoplasmic reticulum (ER), and Calr3b, a Ca^2+^ dependent ER chaperone were increased, along with ECM and metabolism-related proteins (**Supplemental Figure 4** and **Supplemental Tables 6 and 13**). Data from *cryaba^-/-^*; *nrf2^fh318/fh318^* lenses showed changes resembling a combination of both single genotypes (**Supplemental Figure 5** and **Supplemental Tables 7 and 14**).

### Proteomic analysis of lenses 4 days post-PBS vs. H_2_O_2_ treatment

diaPASEF mass spectrometry proteomics revealed that all H_2_O_2_-treated lenses exhibited significant but bidirectional changes in the abundance of cytoskeletal and ECM proteins, with no consistent pattern of increased or decreased protein expression. For instance, different types of collagens were either significantly upregulated or downregulated in the same sample type after H_2_O_2_ treatment. Other protein groups altered by H_2_O_2_ treatment included an increase in energy production, degradation, and protease inhibitor proteins, while ribosomal components and translation associated GO terms were reduced in all H_2_O_2_-treated samples.

Specifically, in WT lenses 4 days post-PBS vs. H_2_O_2_ treatment, Atp2a3 was significantly increased in H_2_O_2_-treated lenses, alongside proteins involved in protein folding, such as Calr3b, Pdia5, and Fkbp9. (**Figure 2** and **Supplemental Tables 8 and 15**). *cryaba^-/-^* lenses 4 days post-PBS vs. H_2_O_2_ treatment also exhibited an increase in protein folding proteins in H_2_O_2_-treated lenses, including Hspa5, Hyou1, Calr3b, Selenof, Hspd1, and Hspe1, alongside Atp2b1a, a plasma membrane Ca^2+^-ATPase. However, there were more significantly changed GO terms in the *cryaba^-/-^* H_2_O_2_-treated lenses compared to the WT H_2_O_2_-treated lenses, including an increase in calcium ion binding, electron transport chain, vesicle organization, and Golgi vesicle transport terms (**Figure 3** and **Supplemental Tables 9 and 16**). In contrast, there were very few significantly differentially abundant proteins in *nrf2^fh318/fh318^* lenses (**Figure 4** and **Supplemental Tables 10 and 17**) after H_2_O_2_ treatment. The *cryaba^-/-^; nrf2^fh318/fh318^* lenses had the largest number of proteins involved in protein folding and calcium homeostasis identified as significantly upregulated in H_2_O_2_-treated lenses, including Hsp90b1, Hyou1, Fkbp9, Calr3b, Hspd1, and Atp2a3. (**Figure 5** and **Supplemental Tables 11 and 18**).

**Figure 2.**
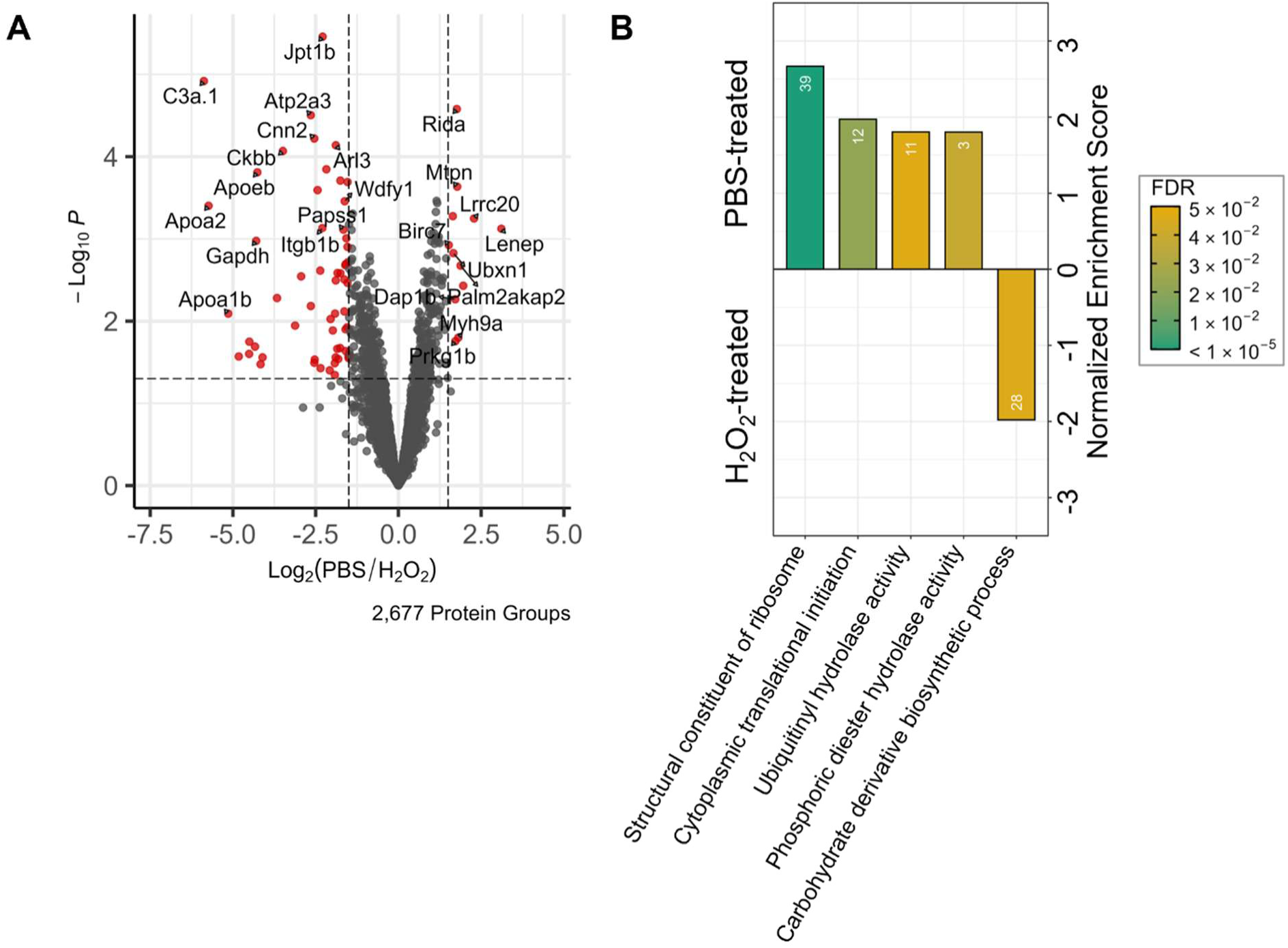
Analysis of WT lenses 4 days post-PBS vs. H_2_O_2_ treatment. **A)** Volcano plot with significant protein groups shown in red with cut-offs of log_2_FC > 1.5 or log_2_FC < -1.5 and p-value < 0.05. **B)** Enriched biological process and molecular function parent GO terms from GSEA ordered by normalized enrichment score. Bars are colored by FDR and only terms with FDR less than or equal to 0.05 are shown.

**Figure 3.**
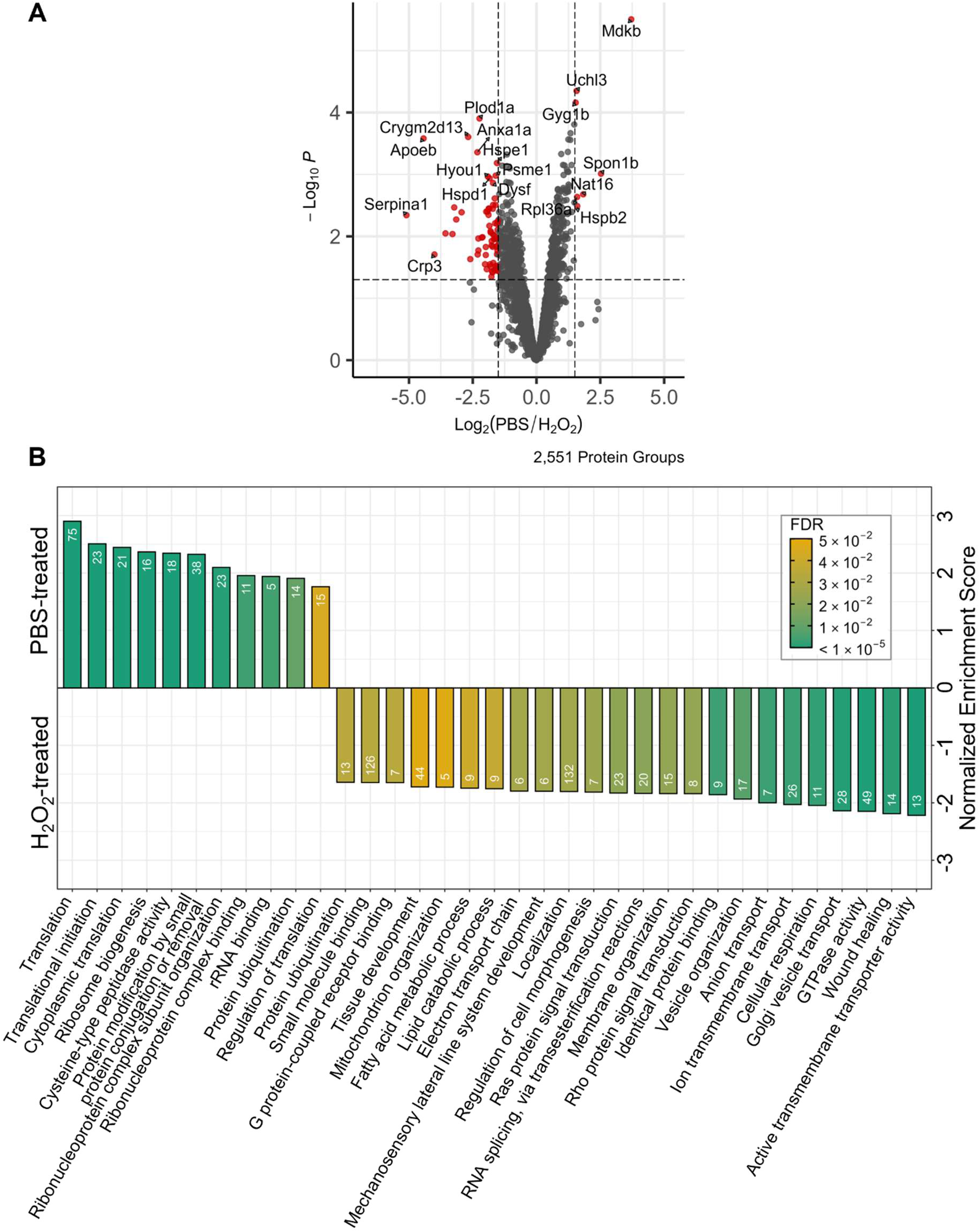
Analysis of *cryaba^-/-^* lenses 4 days post-PBS vs. H_2_O_2_ treatment. **A)** Volcano plot with significant protein groups shown in red with cut-offs of log_2_FC > 1.5 or log_2_FC < -1.5 and p-value < 0.05. **B)** Enriched biological process and molecular function parent GO terms from GSEA ordered by normalized enrichment score. Bars are colored by FDR and only terms with FDR less than or equal to 0.05 are shown.

**Figure 4.**
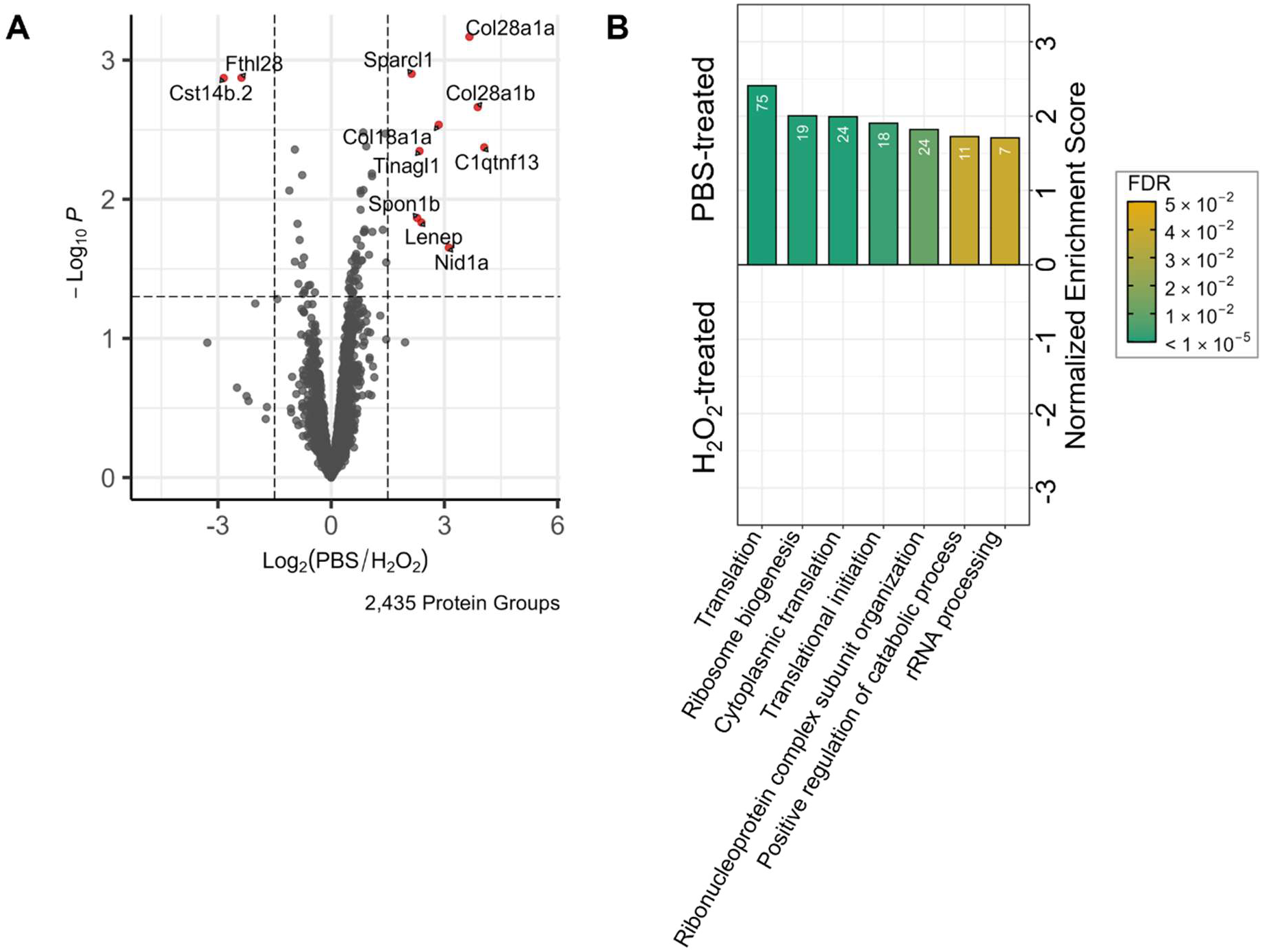
Analysis of *nrf2^fh318/fh318^* lenses 4 days post-PBS vs. H_2_O_2_ treatment. **A)** Volcano plot with significant protein groups shown in red with cut-offs of log_2_FC > 1.5 or log_2_FC < -1.5 and p-value < 0.05. **B)** Enriched biological process and molecular function parent GO terms from GSEA ordered by normalized enrichment score. Bars are colored by FDR and only terms with FDR less than or equal to 0.05 are shown.

**Figure 5.**
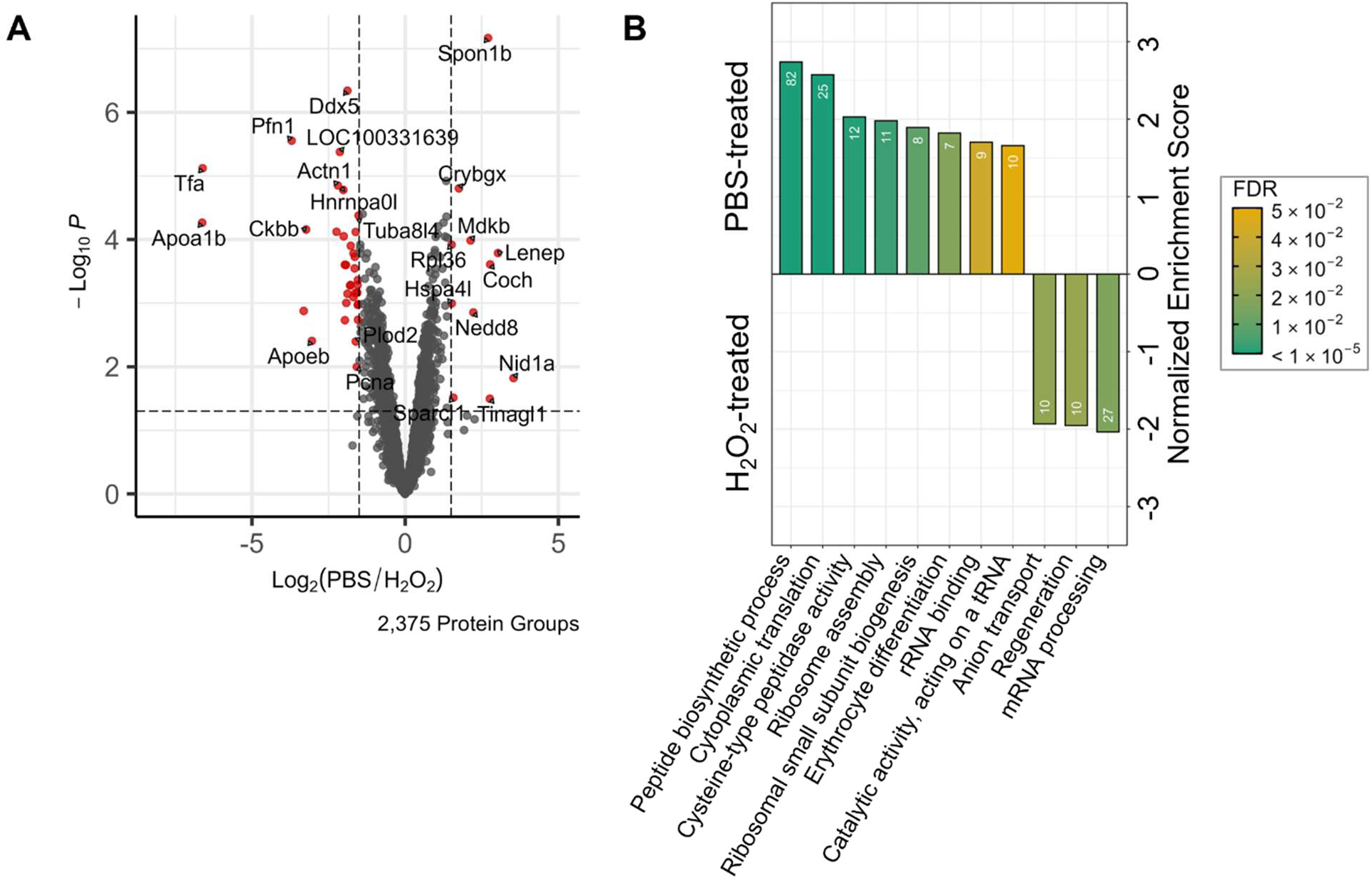
Analysis of *cryaba^-/-^*; *nrf2^fh318/fh318^* lenses 4 days post-PBS vs. H_2_O_2_ treatment. **A)** Volcano plot with significant protein groups shown in red with cut-offs of log_2_FC > 1.5 or log_2_FC < -1.5 and p-value < 0.05. **B)** Enriched biological process and molecular function parent GO terms from GSEA ordered by normalized enrichment score. Bars are colored by FDR and only terms with FDR less than or equal to 0.05 are shown.

## Discussion

While Park et al. identified an interaction between Cryaba and Nrf2 in embryonic zebrafish lenses, the proteomic changes that might be responsible for the observed phenotypes remained unknown.^16^ Furthermore, it also remained unclear how these lenses respond to oxidative stress in the adult zebrafish. Therefore, zebrafish lenses were treated with PBS or H_2_O_2_, resulting opacities were assessed, and lenses underwent proteomic analysis.

The significant increase in opacity in the WT vs. *cryaba^-/-^* lenses for all days and treatments pooled together in the adult zebrafish is similar to Park et al., who reported that the lenses of *cryaba^-/-^* embryos had significantly more defects compared to the lenses of WT embryos.^16^ However, Park et al. only examined untreated embryos. Additionally, the significant increase in opacity of *nrf2^fh318/fh318^* lenses compared to WT lenses for all days and treatments pooled together may be due to the accumulation of oxidative stress-induced damage during the zebrafish lens aging process. The lack of a significant treatment and day interaction in the ANOVA opacity results suggest that a timepoint beyond 8 days post-injection is needed to observe cataract resolution.

### Mutant lenses 4 days post-PBS treatment may be unable to properly differentiate

In the *cryaba^-/-^* zebrafish, the decrease in cytoskeletal and integral membrane and membrane-associated proteins such as Aqp0a and Gja8 suggests reduced membrane stability, cell adhesion, and impacts to the lens microcirculatory system.^32^ In the lens, epithelial cells elongate and differentiate into organelle-free lens fiber cells that compose the cortex and nucleus regions.^1,3,4^ Together, these changes support a shift away from normal differentiation, as lens fiber cells may fail to properly elongate and exhibit abnormal packing, an unreported phenotype. Furthermore, the increase in metabolism, catabolism, and translation-associated proteins suggests that organelles may not be properly degrading during differentiation. Similarly, the *nrf2^fh318/fh318^* lenses also exhibit a shift away from normal differentiation as evidenced by an increase in metabolism and ECM proteins, while the decrease in Calb1 and the increase in Atp2a3 are consistent with the established role of Nrf2 in calcium homeostasis. *cryaba^-/-^*; *nrf2^fh318/fh318^* lenses exhibited a similar pattern.^33^ Significantly altered pathways for each mutant comparison are summarized in **Table 1**.

**Table 1.** Significantly altered pathways in WT vs. mutant lenses 4 days post-PBS treatment.

| Genotype | Significant pathways when compared to WT |
| --- | --- |
| *cryaba*<sup>-/-</sup> | **Unable to properly differentiate**<br>(↓Cryaba, Pkp3b, Aqp0a, Gja8, Cadm2a) |
| *nrf2*<sup>fh318/fh318</sup> | **Unable to properly differentiate** (↑Impdh1b, Gapdh, Hspg2),<br>**dysregulated calcium homeostasis** (↓Calb1, ↑Atp2a3) |
| *cryaba*<sup>-/-</sup>;<br>*nrf2*<sup>fh318/fh318</sup> | **Unable to properly differentiate** (↓Cryaba, Crygm2d12, Sdk2a, ↑Impdh1b, Gapdh), **dysregulated calcium homeostasis** (↑Icn) |

### Lenses 4 days post-H_2_O_2_ treatment exhibit upregulation of chaperones and the unfolded protein response

In WT lenses 4 days post-H_2_O_2_ treatment, a significant increase in the ER-resident chaperones Calr3b, Pdia5, Fkbp9, and ER Ca^2+^ pump Atp2a3, indicates the activation of the unfolded protein response (UPR). H_2_O_2_ oxidizes ER membrane Ca^2+^ release channels, allowing Ca^2+^ from the ER to leak into the cytosol.^34^ Elevated cytosolic Ca^2+^ levels can then activate calpains, which cleave proteins and cause protein aggregation. The activation of calpains in rats via selenite injections results in the formation of cataracts.^35–37^ The depletion of ER Ca^2+^ stores perturbs the function of ER chaperones leading to an accumulation of misfolded proteins. Hspa5 then dissociates from the UPR sensors Perk, Ire1, and Atf6 to activate the UPR.^38^ Translation is also globally reduced to diminish overload of misfolded proteins, which is consistent with the observed statistically significant decrease in ribosomal proteins and translation-associated GO terms in all H_2_O_2_-treated lenses. The uniform decrease in translation in all H_2_O_2_-treated lenses also signals a significant change in the differentiation of lens fiber cells. However, the expression of resident-ER chaperones and folding enzymes are selectively increased in the UPR.^39^ The increase in the ER Ca^2+^-ATPase Atp2a3 suggests a compensatory attempt to restore depleted ER Ca^2+^ stores.

*cryaba^-/-^*lenses also exhibit the activation of the UPR 4 days post-H_2_O_2_ treatment, while the significantly increased expression of Atp2b1a reflects an overwhelmed capacity to remove Ca^2+^ from the cytosol. The increased expression of ATP-dependent mitochondrial matrix chaperonins Hspd1 and Hspe1 in *cryaba^-/-^* lenses suggests that the mitochondrial unfolded protein response (UPR^mt^) is also activated. The UPR^mt^ is activated when mitochondria are exposed to stress, allowing Atf5 to translocate to the nucleus and upregulate the transcription of stress-response genes.^40^ The combined activation of the UPR and UPR^mt^ indicates that the *cryaba^-/-^* lenses 4 days post-H_2_O_2_ treatment are responding to stress in multiple cellular compartments.

The complete absence of significantly increased chaperones in *nrf2^fh318/fh318^* lenses 4 days post-H_2_O_2_ treatment suggests UPR signaling was initiated, as translation was still suppressed, but could not be sustained. While Nrf2 is not known to directly control chaperone transcription, Nrf2 loss of function impairs antioxidant gene expression. Thus, Nrf2 loss of function may leave the ER overwhelmed with misfolded proteins, preventing the sustained activation of the UPR.^41^

The *cryaba^-/-^*; *nrf2^fh318/fh318^* H_2_O_2_-treated lenses exhibit chaperone expression patterns matching H_2_O_2_-treated *cryaba^-/-^* lenses. Specifically, the UPR and UPR^mt^ pathways are activated, and the increase in Atp2a3 also indicates that *cryaba^-/-^*; *nrf2^fh318/fh318^* lenses 4 days post-H_2_O_2_ treatment may exhibit similar Ca^2+^ levels as WT lenses post-H_2_O_2_ treatment. However, unlike the *nrf2^fh318/fh318^* 4 days post-H_2_O_2_ treatment lenses, the activation of the UPR is sustained despite loss of Nrf2 function, suggesting that the activation of the UPR in this case may be Nrf2- independent. Significantly altered UPR and UPR^mt^ pathways in each zebrafish line after H_2_O_2_ treatment are summarized in **Table 2**.

**Table 2.** Significantly altered UPR and UPR^mt^ pathways in all zebrafish lenses 4 days post-H_2_O_2_ treatment.

| Genotype | Significant pathways |
| --- | --- |
| WT | UPR (↑Calr3b, Pdia5, Fkbp9) |
| *cryaba*<sup>-/-</sup> | UPR (↑Hspa5, Hyou1, Calr3b, Selenof), UPR<sup>mt</sup> (↑Hspd1, Hspe1) |
| *nrf2*<sup>fh318/fh318</sup> | UPR (Previously activated but no longer sustained by 4 days post-H<sub>2</sub>O<sub>2</sub> treatment) |
| *cryaba*<sup>-/-</sup>; *nrf2*<sup>fh318/fh318</sup> | UPR (↑Hsp90b1, Hyou1, Fkbp9, Calr3b), UPR<sup>mt</sup> (↑Hspd1) |

### Other pathways upregulated in H_2_O_2_-treated lenses

Activation of the UPR also activates the ER-associated degradation (ERAD) pathway and lysosomal proteolysis-related proteins to further reduce ER-associated stress and the removal of misfolded proteins.^42^ Dnajb11, an ERAD-associated co-chaperone, was detected only in WT H_2_O_2_-treated lenses and not in WT PBS-treated lenses.^43^ The increased expression of protease inhibitors in H_2_O_2_-treated lenses also indicates robust control of proteolysis during oxidative stress to prevent excessive protein degradation, while. the increase in ATP-producing proteins may reflect the increase in energy needed to sustain the UPR.^39,44^ Lastly, the observed difficulty in the separation of H_2_O_2_-treated lenses may reflect H_2_O_2_-induced cytoskeletal and ECM remodeling via the simultaneous increase and decrease in cytoskeletal and ECM proteins, which could alter the mechanical properties of the lens tissue.

### Comparison of results to other datasets

Cholesterol is highly abundant in lens fiber cell membranes and may play a protective role against ARC formation.^45–47^ Park et al. demonstrated the cholesterol biosynthesis pathway was transcriptionally upregulated in adult *cryaba^-/-^*; *nrf2^fh318/fh318^* lenses.^16^ However, no corresponding cholesterol biosynthesis proteins were detected in the present dataset. Future proteomic studies using membrane-enriched samples may validate whether the cholesterol biosynthesis pathway is activated to help prevent oxidative-stress induced cataract formation.^48^

Data-independent acquisition mass spectrometry proteomics has been used to analyze the phacoemulsified lens contents from ARC patients vs. clear lenses.^49^ In ARC samples, cytoskeletal proteins were significantly downregulated, while in the H_2_O_2_-injected zebrafish model, significant bidirectional changes in cytoskeletal and ECM remodeling pathways were identified. This distinct pattern in the zebrafish model likely reflects an early acute stress response that precedes the strong downregulation observed in ARC lenses, indicating that analysis of both sample types is needed to fully understand cataract pathogenesis.

The lens transcriptome of WT zebrafish 0, 14, and 30 days post-H_2_O_2_ injection has previously been examined, with significant changes focused on miRNA.^18,20^ Combined with the changes observed in this dataset, these findings suggest a phased response to oxidative stress in the zebrafish lens, in which the activation of the UPR and UPR^mt^ at the protein level may precede miRNA-driven changes.

The activation of the UPR has previously been observed in healthy aged and ARC lenses.^50–52^ However, the diaPASEF mass spectrometry proteomics approach extends these findings by revealing temporal and genotype-specific UPR and UPR^mt^ dynamics and changes to other related pathways during cataract formation (**Figure 6**). The observation that H_2_O_2_-treated zebrafish lenses recapitulate pathways previously identified in aged and ARC lenses further support oxidative stress as a key driver of ARC development.

**Figure 6.**
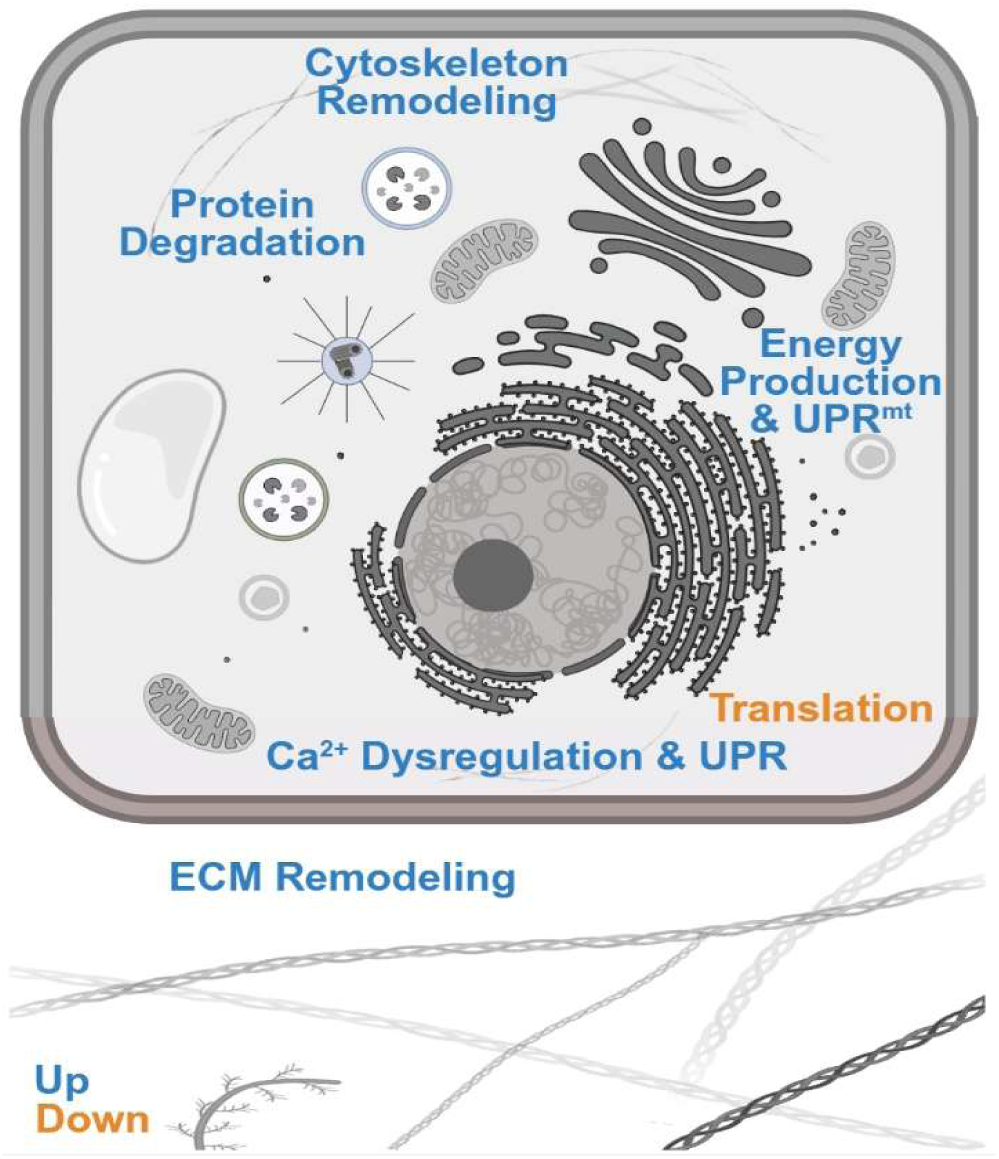
Diagram of pathways that were significantly changed following H_2_O_2_-treatment in the lens. Created in BioRender. Zelle, S. (2026) https://BioRender.com/bs07i6l.

## Conclusion

The data demonstrated that treatment with H_2_O_2_ resulted in the formation of cortical cataracts in the zebrafish lens. Proteomic findings revealed that H_2_O_2_-induced oxidative stress, dependent on genotype, triggers the UPR and UPR^mt^ in zebrafish lenses 4 days post-treatment, confirming critical interactions between Cryaba and Nrf2 in the lens. Future post-translational modification (PTM)-specific proteomics studies are needed to better understand which proteins in the lens become modified with oxidative PTMs, as the present dataset only focused on changes in protein abundance.

While this study captured strong treatment-driven changes in the lens proteome, the high biological variability found in zebrafish suggests that additional studies with increased biological replicates may further resolve subtle changes in the proteome that did not pass the significance threshold. Together, the pathways identified in this dataset represent a framework for understanding the molecular mechanisms that link oxidative stress to ARC formation. Key proteins within these pathways represent potential therapeutic targets for the development of drugs aimed at delaying or inhibiting ARC formation.

## Author Contributions

S.R.Z., H.S.M., and K.L.S.: conceptualization, S.R.Z.: sample preparation, W.H.M.: data acquisition, H.S.M. and K.L.S.: resources and supervision, S.R.Z.: data analysis, S.R.Z.: writing-original draft, S.R.Z., W.H.M., H.S.M., and K.L.S.: writing-review & editing, S.R.Z., H.S.M., and K.L.S.: funding acquisition.

## Acknowledgments

The authors thank Zhen Wang and Lee Cantrell for discussions of this project and Wenbiao Chen for providing zebrafish expertise and equipment. The authors also acknowledge support from the Vanderbilt Mass Spectrometry Research Center, particularly Kristie Rose and Purvi Patel, and Jenny Schafer and Oleg Kovtun of the Vanderbilt Cell Imaging Shared Resource, supported by NIH grants CA68485, DK20593, DK58404, DK59637 and EY08126. ChatGPT, specifically models GPT 4o, GPT 4o mini, GPT 4.1 mini, o4 mini, GPT 5, and GPT 5 mini was used to assist with the generation of some blocks of R code in mid-2025. All suggestions were manually reviewed, and the authors maintain responsibility for all changes.

