## Supplemental File 1 for "Upregulation of the Unfolded and Mitochondrial Unfolded Protein Responses in Oxidative Stress-Induced Cataract"

// select input and output directories

inputDir = getDirectory("Select a folder containing .czi images");

outputDir = getDirectory("Select a folder to save results");

// set pixel-to-micron scale

micron_per_pixel = 1.1364;

bin_size = 10 / micron_per_pixel;

min_area_microns = 100000;

min_area_pixels = min_area_microns / (micron_per_pixel * micron_per_pixel);

list = getFileList(inputDir);

for (i = 0; i < list.length; i++) {

filename = list[i];

if (endsWith(filename, ".czi")) {

print("\\Clear");

print("Processing: " + filename);

run("Bio-Formats", "open=[" + inputDir + filename + "] color_mode=Default rois_import=[ROI manager] view=Hyperstack stack_order=XYCZT");

baseName = replace(filename, ".czi", "");

run("RGB Color");

run("8-bit");

rename("original");

run("Duplicate...", "title=mask");

run("Enhance Contrast...", "saturated=0.35 normalize equalize");

run("Enhance Local Contrast (CLAHE)", "blocksize=127 histogram=256 maximum=3 mask=*None* fast_(less_accurate)");

run("Median...", "radius=4");

setAutoThreshold("Default dark");

//run("Threshold...");

setThreshold(75, 255, "raw");

//setThreshold(75, 255);

setOption("BlackBackground", true);

run("Convert to Mask");

run("Fill Holes");

run("Watershed");

run("Analyze Particles...", "size=" + min_area_pixels + "-Infinity show=Masks clear include add");

run("Distance Map");

run("Enhance Contrast", "saturated=0.35");

run("Invert");

run("Find Maxima...", "prominence=0 light output=[List]");

// read detected maxima coordinates from the Results Table

x_coords = Table.getColumn("X");

y_coords = Table.getColumn("Y");

num_maxima = x_coords.length;

// print headers in the log window

print("ROI,Radius (microns),Mean Intensity (in annular ring)");

// set up a for loop for each center if you two eyes present

for (roi_index = 0; roi_index < num_maxima; roi_index++) {

x = x_coords[roi_index];

y = y_coords[roi_index];

// get themax radius for current ROI using the distance transform image

selectWindow("Mask of mask");

roiManager("Select", roi_index);

run("Measure");

// Extract the max value from the distance transform within the ROI

getStatistics(area, mean, min, max);

max_radius = (sqrt(area / PI))/micron_per_pixel;

// compute mean intensity for each 10 micron thick bin propagating outward from the center

for (r = bin_size; r <= max_radius; r += bin_size) {

// create outer and inner circles

selectWindow("original");

makeOval(x - r, y - r, 2 * r, 2 * r);

run("Measure");

outer_intensity = getResult("Mean", nResults() - 1);

outer_area = getResult("Area", nResults() - 1);

makeOval(x - (r - bin_size), y - (r - bin_size), 2 * (r - bin_size), 2 * (r - bin_size));

run("Measure");

inner_intensity = getResult("Mean", nResults() - 1);

inner_area = getResult("Area", nResults() - 1);

// get mean intensity in the annular ring (the band)

annular_intensity = ((outer_intensity * outer_area) - (inner_intensity * inner_area)) / (outer_area - inner_area);

// print results for current roi into the log window

print(roi_index + 1 + "," + (r * micron_per_pixel) + "," + annular_intensity);

}

}

savePath = outputDir + baseName + "_intensity_profile.csv";

selectWindow("Log");

saveAs("Text", savePath);

selectWindow("Mask of mask");

saveAs("Tiff", outputDir + baseName + "_distance_mask.tif");

roiFilePath = outputDir + baseName + "_ROIs.zip";

roiManager("Deselect");

roiManager("Save", roiFilePath);

}

// Close extra windows; i commented out the line below to be able to visualize the results

run("Close All");

}
