## Supplemental File 5 for "Upregulation of the Unfolded and Mitochondrial Unfolded Protein Responses in Oxidative Stress-Induced Cataract"

**Supplemental Table 1.** **Results of 3-way ANOVA for gray values with interactions treatment, genotype, and day.** There is a significant main effect of treatment, genotype, and day, but no significant interaction terms.

| **Predictor** | **Sum of squares** | **Degrees of freedom** | **F-value** | **p-value** | **Significance** |
| --- | --- | --- | --- | --- | --- |
| Treatment (H₂O₂ vs. PBS) | 193.518 | 1 | 1521.957 | < 0.001 | *** |
| Genotype (all genotypes) | 1.757 | 3 | 4.607 | 0.004 | ** |
| Day (Day 8 vs. Day 4) | 2.256 | 1 | 17.741 | < 0.001 | *** |
| Treatment × Genotype | 0.550 | 3 | 1.441 | 0.232 | n.s. |
| Treatment × Day | 0.200 | 1 | 1.573 | 0.211 | n.s. |
| Genotype × Day | 0.513 | 3 | 1.344 | 0.261 | n.s. |
| Treatment × Genotype × Day | 0.790 | 3 | 2.070 | 0.105 | n.s. |
| Residuals | 26.320 | 207 |  |  |  |

**Supplemental Table 2. Results of estimated marginal means comparing PBS vs. H_2_O_2_ mean gray values with all day and genotype replicates pooled together.** There is a significant difference between PBS and H_2_O_2_ treatments.

| **Contrast** | **Estimate** | **SE** | **t-value** | **p-value** | **Significance** |
| --- | --- | --- | --- | --- | --- |
| H_2_O_2_ vs. PBS | -1.863 | 0.048 | -39.012 | < 0.001 | *** |

**Supplemental Table 3. Results of estimated marginal means comparing mean gray values for each day with all genotype and treatment replicates pooled together.** There is a significant difference between day 4 and 8.

| **Contrast** | **Estimate** | **SE** | **t-value** | **p-value** | **Significance** |
| --- | --- | --- | --- | --- | --- |
| Day 4 vs. Day 8 | 0.201 | 0.0478 | 4.212 | < 0.001 | *** |

**Supplemental Table 4.** **Results of estimated marginal means comparing mean gray values for each genotype with all day and treatment replicates pooled together.** There is a significant difference between WT vs. *cryaba^-/-^* and WT vs. *nrf2^fh318/fh318^*.

| **Contrast** | **Estimate** | **SE** | **t-value** | **p-value** | **Significance** |
| --- | --- | --- | --- | --- | --- |
| WT vs. *cryaba^-/-^* | -0.242 | 0.068 | -3.573 | 0.002 | ** |
| WT vs. *nrf2^fh318/fh318^* | -0.180 | 0.067 | -2.669 | 0.041 | * |
| WT vs. *cryaba^-/-^; nrf2^fh318/fh318^* | -0.137 | 0.067 | -2.039 | 0.177 | n.s. |
| *cryaba^-/-^* vs. *nrf2^fh318/fh318^* | 0.062 | 0.068 | 0.916 | 0.796 | n.s. |
| *cryaba^-/-^* vs. *cryaba^-/-^; nrf2^fh318/fh318^* | 0.104 | 0.068 | 1.543 | 0.414 | n.s. |
| *cryaba^-/-^; nrf2^fh318/fh318^* vs. *nrf2^fh318/fh318^* | 0.042 | 0.067 | 0.630 | 0.922 | n.s. |
