## Supplemental File 6 for "Upregulation of the Unfolded and Mitochondrial Unfolded Protein Responses in Oxidative Stress-Induced Cataract"

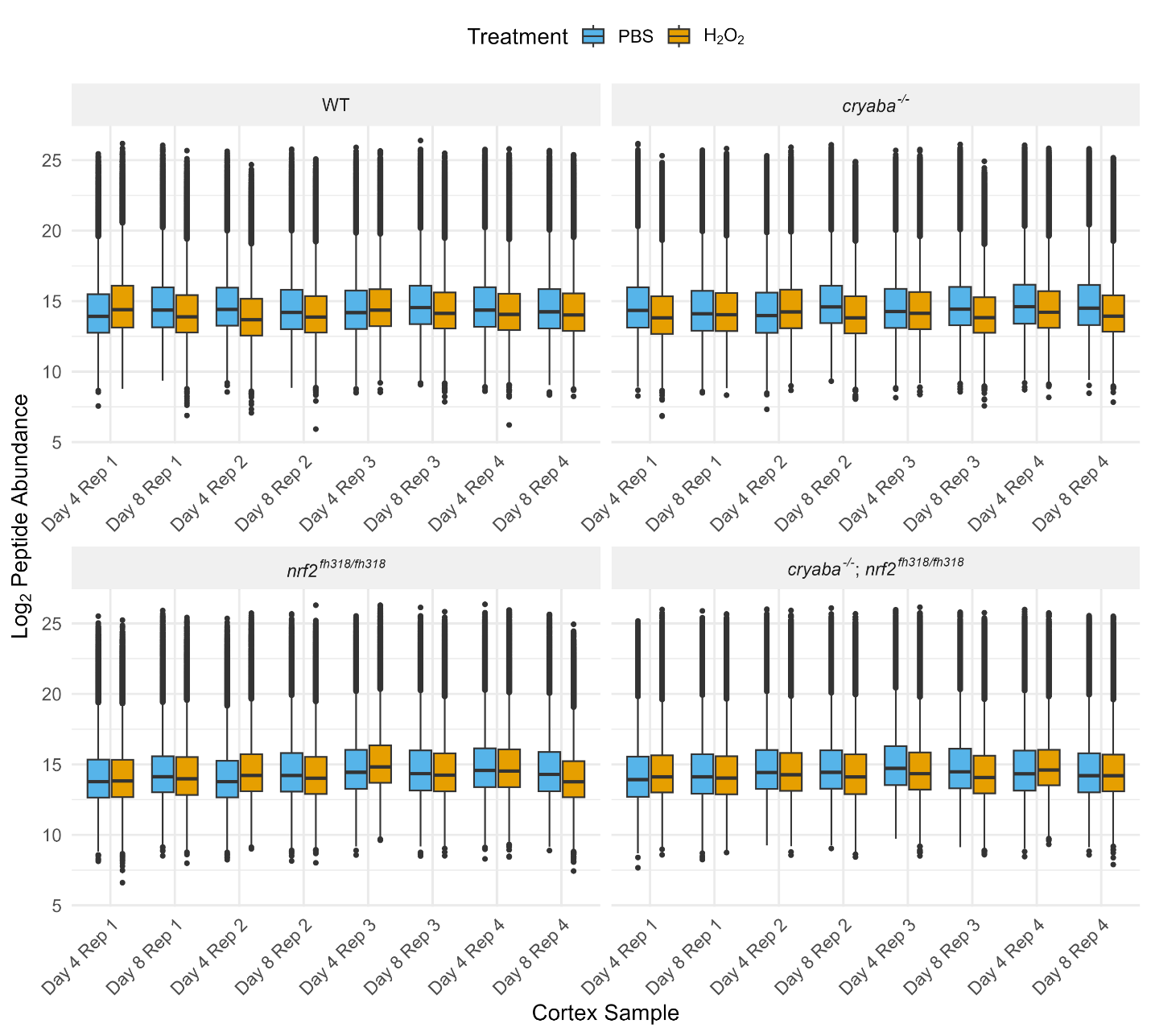

**Supplemental 1. Log_2_ quantity of peptide MS2 intensities after median fold change normalization.**

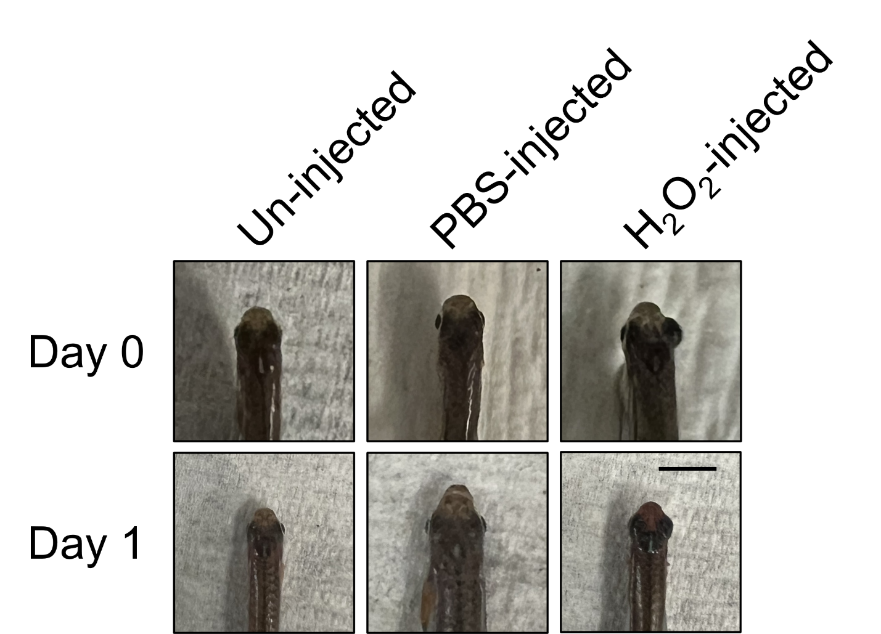

**Supplemental Figure 2. Representative images of un-injected, PBS-injected, and H_2_O_2_-injected eyes 0 days and 1 day post-injection.** Only the right eye was injected. Immediately after successful H_2_O_2_-injection, the eye becomes enlarged, returning to normal by 1 day post-injection. The black scalebar corresponds to 1 cm. The temporary enlargement of the eye observed only in H_2_O_2_-injected eyes may reflect the formation and resolution of a corneal edema via the oxidation of Na^+^/K^+^-ATPase pumps and lipid peroxidation in the corneal endothelium.

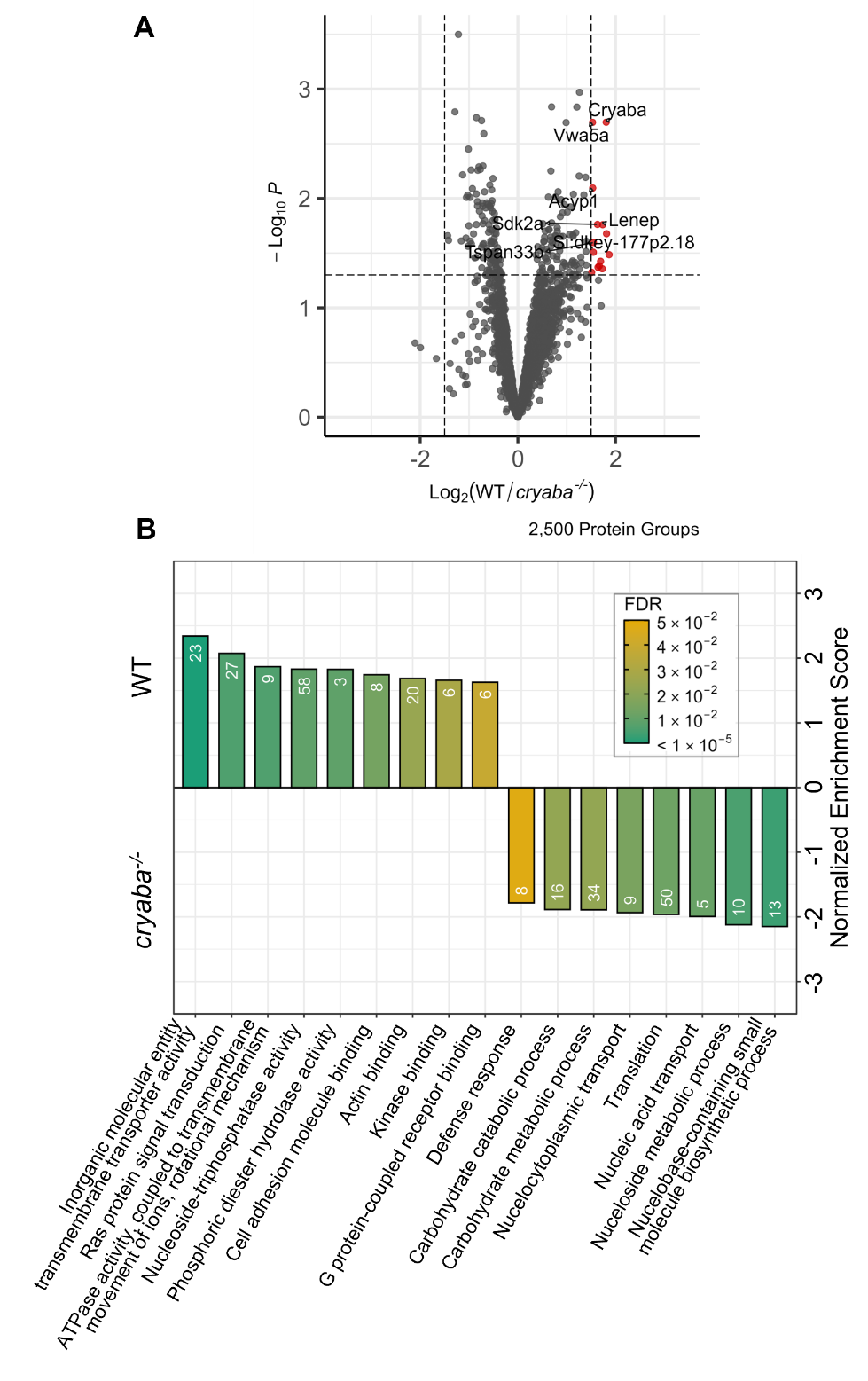

**Supplemental Figure 3. Analysis of WT vs. cryaba^-/-^ lenses 4 days post-PBS treatment. A)** Volcano plot with significant protein groups shown in red with cut-offs of log_2_FC > 1.5 or log_2_FC < -1.5 and p-value < 0.05. **B)** Enriched biological process and molecular function parent GO terms from GSEA ordered by normalized enrichment score. Bars are colored by FDR and only terms with FDR less than or equal to 0.05 are shown.

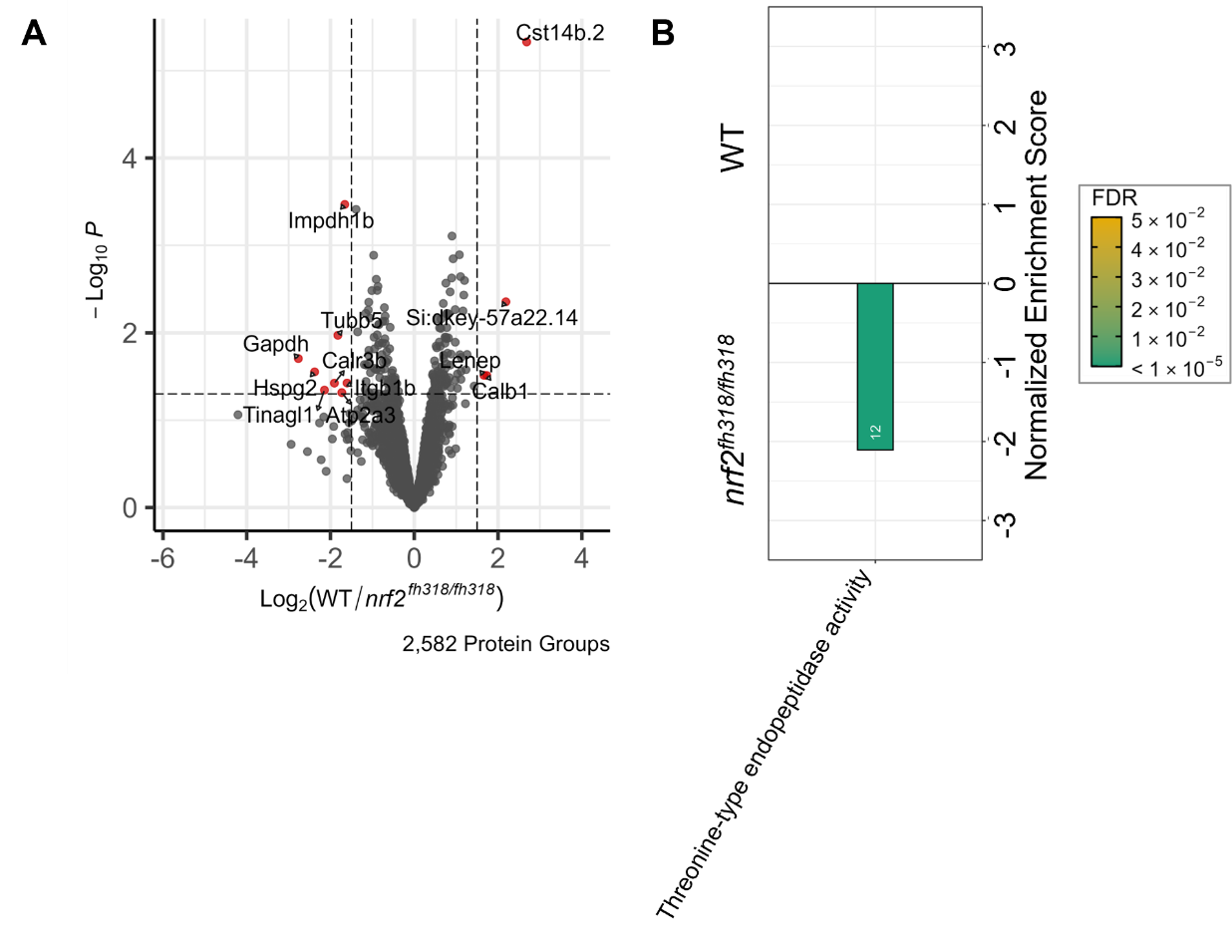

**Supplemental Figure 4. Analysis of WT vs. nrf2^fh318/fh318^ lenses 4 days post-PBS treatment. A)** Volcano plot with significant protein groups shown in red with cut-offs of log_2_FC > 1.5 or log_2_FC < -1.5 and p-value < 0.05. **B)** Enriched biological process and molecular function parent GO terms from GSEA ordered by normalized enrichment score. Bars are colored by FDR and only terms with FDR less than or equal to 0.05 are shown.

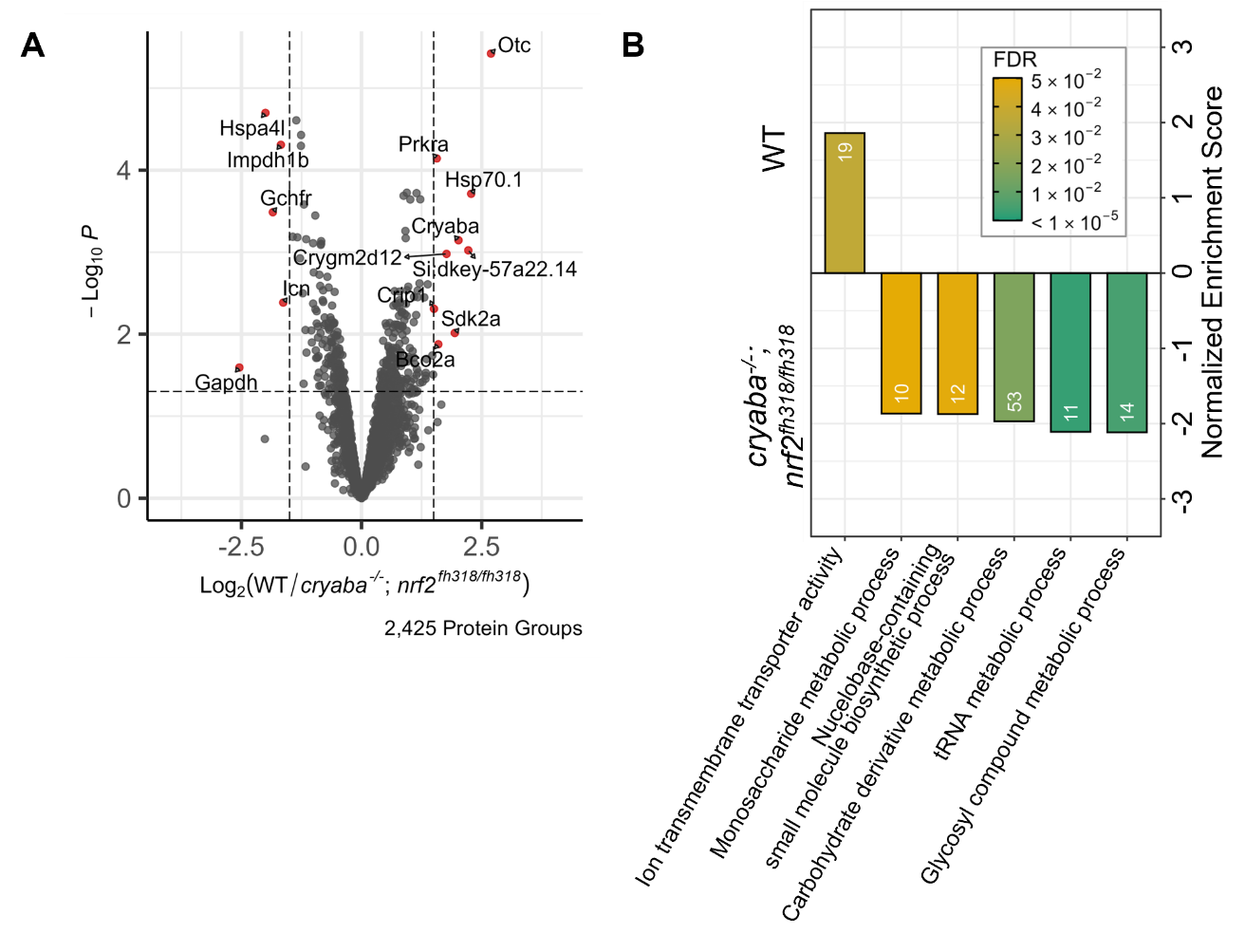

**Supplemental Figure 5. Analysis of WT vs. cryaba^-/-^; nrf2^fh318/fh318^ lenses 4 days post-PBS treatment. A)** Volcano plot with significant protein groups shown in red with cut-offs of log_2_FC > 1.5 or log_2_FC < -1.5 and p-value < 0.05. **B)** Enriched biological process and molecular function parent GO terms from GSEA ordered by normalized enrichment score. Bars are colored by FDR and only terms with FDR less than or equal to 0.05 are shown.
