## Supplemental File 7 for "Upregulation of the Unfolded and Mitochondrial Unfolded Protein Responses in Oxidative Stress-Induced Cataract"

**Supplemental Table 5. Significant proteins (p-value < 0.05 and log_2_FC > 1.5 or log_2_FC < -1.5) from WT 4 days post-PBS treated lenses compared to *cryaba^-/-^* 4 days post-PBS treated lenses organized by log_2_FC (n = 14).**

| **Gene group** | **Protein group** | **Protein description** | **p-value** | **Log_2_FC** |
| --- | --- | --- | --- | --- |
| *pkp3b* | F1QWE6 | Plakophilin 3b | 0.033 | 1.868 |
| *si:dkey-177p2.18* | A0A8M9PTB0 | Phospholipase B1, membrane-associated | 0.021 | 1.815 |
| *cryaba* | Q9PUR2 | Alpha-crystallin B chain | 0.002 | 1.808 |
| *lenep* | A0A8M1P644 | Lens epithelial cell protein LEP503 | 0.017 | 1.736 |
| *cxadr* | Q90Y50 | Coxsackievirus and adenovirus receptor homolog | 0.044 | 1.728 |
| *nrcamb* | A0A8M9PV37 | Neuronal cell adhesion molecule | 0.038 | 1.693 |
| *gja8a* | A0A8N1Z0D3 | Gap junction protein | 0.041 | 1.668 |
| *lctlb* | A0A286Y8V0 | Lactase-like b | 0.043 | 1.64 |
| *sdk2a* | A0AB32TJL9 | Protein sidekick-2 isoform X7 | 0.017 | 1.63 |
| *tspan33b* | A0A8M2B5Z3 | Tetraspanin | 0.025 | 1.548 |
| *aqp0a* | Q6DEI6 | Aquaporin-0a | 0.031 | 1.545 |
| *acyp1* | A3KPA7 | Acylphosphatase | 0.008 | 1.532 |
| *vwa5a* | Q66HV5 | Zgc:92481 | 0.002 | 1.53 |
| *cadm2a* | A0A8M2B1M0 | Cell adhesion molecule 2a isoform X1 | 0.047 | 1.51 |

**Supplemental Table 6. Significant proteins (p-value < 0.05 and log_2_FC > 1.5 or log_2_FC < -1.5) from WT 4 days post-PBS treated lenses compared to *nrf2^fh318/fh318^* 4 days post-PBS treated lenses organized by log_2_FC (n = 12).**

| **Gene group** | **Protein group** | **Protein description** | **p-value** | **Log_2_FC** |
| --- | --- | --- | --- | --- |
| *cst14b.2* | A0A2R8RU89 | Cystatin 14b, tandem duplicate 2 | < 0.001 | 2.684 |
| *si:dkey-57a22.14* | A0AB32TXN5 | Gamma-crystallin D | 0.004 | 2.188 |
| *calb1* | E9QJF8 | Calbindin | 0.031 | 1.732 |
| *lenep* | A0A8M1P644 | Lens epithelial cell protein LEP503 | 0.031 | 1.664 |
| *itgb1b* | Q3YAA0 | Integrin beta | 0.038 | -1.605 |
| *impdh1b* | Q5RGV1 | Inosine-5'-monophosphate dehydrogenase 1b | < 0.001 | -1.66 |
| *atp2a3* | A0A2R8QJH4 | Calcium-transporting ATPase | 0.048 | -1.73 |
| *tubb5* | Q6NW90 | Tubulin beta chain | 0.011 | -1.824 |
| *calr3b* | A0A0R4IL29 | Calreticulin | 0.038 | -1.907 |
| *tinagl1* | A3KNG7 | Si:dkey-158b13.1 | 0.045 | -2.145 |
| *hspg2* | A0A8M9PWV8 | Basement membrane-specific heparan sulfate proteoglycan core protein isoform X1 | 0.028 | -2.379 |
| *gapdh* | Q5XJ10 | Glyceraldehyde-3-phosphate dehydrogenase | 0.02 | -2.768 |

**Supplemental Table 7. Significant proteins (p-value < 0.05 and log_2_FC > 1.5 or log_2_FC < -1.5) from WT 4 days post-PBS treated lenses compared to *cryaba^-/-^;* *nrf2^fh318/fh318^* 4 days post-PBS treated lenses organized by log_2_FC (n = 14).**

| **Gene group** | **Protein group** | **Protein description** | **p-value** | **Log_2_FC** |
| --- | --- | --- | --- | --- |
| *otc* | E9QHD9 | Ornithine carbamoyltransferase | < 0.001 | 2.694 |
| *hsp70.1* | B0UXR9 | Heat shock cognate 70-kd protein, tandem duplicate 2 | < 0.001 | 2.281 |
| *si:dkey-57a22.14* | A0AB32TXN5 | Gamma-crystallin D | < 0.001 | 2.222 |
| *cryaba* | Q9PUR2 | Alpha-crystallin B chain | < 0.001 | 2.018 |
| *sdk2a* | A0AB32TJL9 | Protein sidekick-2 isoform X7 | 0.010 | 1.941 |
| *crygm2d12* | A7E2K7 | Crystallin, gamma M2d12 | 0.001 | 1.768 |
| *bco2a* | F1QU45 | Carotenoid-cleaving dioxygenase, mitochondrial | 0.013 | 1.598 |
| *prkra* | B0V3F8 | Interferon-inducible double-stranded RNA-dependent protein kinase activator A homolog | < 0.001 | 1.566 |
| *crip1* | Q5BKV7 | Cysteine-rich protein 1 | 0.005 | 1.507 |
| *icn* | Q6XG62 | Protein S100 | 0.004 | -1.631 |
| *impdh1b* | Q5RGV1 | Inosine-5'-monophosphate dehydrogenase 1b | < 0.001 | -1.679 |
| *gchfr* | Q6PBT6 | GTP cyclohydrolase 1 feedback regulatory protein | < 0.001 | -1.847 |
| *hspa4l* | X1WBR5 | Heat shock 70 kDa protein 4L | < 0.001 | -2.001 |
| *gapdh* | Q5XJ10 | Glyceraldehyde-3-phosphate dehydrogenase | 0.026 | -2.544 |

**Supplemental Table 8. Significant proteins (p-value < 0.05 and log_2_FC > 1.5 or log_2_FC < -1.5) from WT 4 days post-PBS treated lenses compared to WT 4 days post-H_2_O_2_ treated lenses organized by log_2_FC (n = 69).**

| **Gene group** | **Protein group** | **Protein description** | **p-value** | **Log_2_FC** |
| --- | --- | --- | --- | --- |
| *lenep* | A0A8M1P644 | Lens epithelial cell protein LEP503 | 0.001 | 3.110 |
| *lrrc20* | A0A8M9QMF3 | Leucine-rich repeat-containing protein 20 isoform X1 | 0.001 | 2.284 |
| *si:dkey-57a22.14* | A0AB32TXN5 | Gamma-crystallin D | 0.004 | 1.957 |
| *palm2akap2* | A0AB32U0R7 | Palm2 and akap2 fusion isoform X1 | 0.002 | 1.876 |
| *myh9a* | A0A8M1NEM1 | Myosin-9 | 0.016 | 1.810 |
| *mtpn* | Q7T2B9 | Myotrophin | < 0.001 | 1.775 |
| *rida* | Q6AXL2 | 2-iminobutanoate/2-iminopropanoate deaminase isoform 1 | < 0.001 | 1.764 |
| *prkg1b* | A0A8M9Q0L9 | CGMP-dependent protein kinase 1-like | 0.018 | 1.721 |
| *dap1b* | Q9I9N0 | Death-associated protein-like 1 homolog | 0.005 | 1.716 |
| *ubxn1* | Q6NXA9 | UBX domain-containing protein 1 | 0.001 | 1.661 |
| *si:ch211-59p23.1* | A0AB13A846 | Adipogenesis regulatory factor protein-like | 0.001 | 1.643 |
| *birc7* | A0AB13AAI8 | Baculoviral IAP repeat-containing protein 7 isoform 1 | 0.001 | 1.518 |
| *tpm3* | A0A2R8Q650 | Tropomyosin 3 | 0.028 | -1.505 |
| *ywhaqa* | Q7ZUM0 | Tyrosine 3-monooxygenase/tryptophan 5-monooxygenase activation protein, theta polypeptide a | 0.027 | -1.509 |
| *gale* | F1Q5H4 | UDP-glucose 4epimerase | 0.002 | -1.525 |
| *pdia5* | A8WG75 | Pdia5 protein | < 0.001 | -1.538 |
| *pfkpb* | A0AB32THG1 | ATP-dependent 6-phosphofructokinase, platelet type isoform X2 | 0.001 | -1.538 |
| *etfb* | Q7ZV76 | Electron transfer flavoprotein subunit beta | 0.003 | -1.538 |
| *ace* | E7FFA5 | Angiotensin-converting enzyme | 0.012 | -1.550 |
| *nop58* | Q6P6X6 | NOP58 ribonucleoprotein homolog (yeast) | 0.001 | -1.573 |
| *nt5c1bb* | F1QI68 | 5'-nucleotidase, cytosolic IB b | 0.002 | -1.591 |
| *plod1a* | A8WIN7 | Procollagen-lysine,2-oxoglutarate 5-dioxygenase 1 | 0.023 | -1.591 |
| *dkc1* | F1Q749 | H/ACA ribonucleoprotein complex subunit DKC1 | 0.013 | -1.598 |
| *lrpap1* | A0A8M1PJ59 | Alpha-2-macroglobulin receptor-associated protein precursor | 0.002 | -1.604 |
| *wdfy1* | Q4VBV0 | WD repeat and FYVE domain-containing 1 | < 0.001 | -1.623 |
| *mtdha* | A0A8M6YWB0 | Metadherin a isoform X1 | 0.003 | -1.633 |
| *gchfr* | Q6PBT6 | GTP cyclohydrolase 1 feedback regulatory protein | 0.008 | -1.635 |
| *papss1* | B3DJF3 | 3'-phosphoadenosine 5'-phosphosulfate synthase 1 | 0.001 | -1.661 |
| *acta1b* | Q4KMI7 | Actin alpha 1, skeletal muscle b | 0.003 | -1.752 |
| *dnajc10* | A4IG47 | DnaJ homolog subfamily C member 10 | < 0.001 | -1.754 |
| *vclb* | A0A0S2I7K2 | Vinculin | 0.021 | -1.760 |
| *csrp1a* | Q6NZV4 | Cysteine and glycine-rich protein 1 | 0.029 | -1.819 |
| *slc25a24* | Q66L49 | Mitochondrial adenyl nucleotide antiporter SLC25A24 | 0.003 | -1.839 |
| *plod2* | A4U7F9 | Procollagen-lysine 5-dioxygenase | 0.022 | -1.847 |
| *arl3* | Q1MTE5 | ADP-ribosylation factor-like protein 3 | < 0.001 | -1.890 |
| *impdh1b* | Q5RGV1 | Inosine-5'-monophosphate dehydrogenase 1b | 0.003 | -1.892 |
| *tubb5* | Q6NW90 | Tubulin beta chain | 0.028 | -1.895 |
| *tln1* | A0A8M2B1K5 | Talin-1 isoform X1 | 0.008 | -1.914 |
| *pcna* | Q9PTP1 | Proliferating cell nuclear antigen | 0.045 | -1.922 |
| *hnrnpa0l* | Q7ZU48 | Heterogeneous nuclear ribonucleoprotein A0,-like | 0.032 | -1.922 |
| *fkbp9* | Q6DBZ1 | Peptidylprolyl isomerase | 0.013 | -1.983 |
| *myh9b* | A0A8M1QUS6 | Myosin-9 | 0.009 | -2.050 |
| *LOC100331639* | A0A8M1RDD4 | Myosin heavy chain, embryonic smooth muscle isoform | 0.040 | -2.075 |
| *tpm4a* | F1R412 | Tropomyosin 4a | < 0.001 | -2.176 |
| *jpt1b* | A0A8M2BAY7 | Jupiter microtubule associated homolog 1b isoform X1 | < 0.001 | -2.289 |
| *itgb1b* | Q3YAA0 | Integrin beta | 0.001 | -2.298 |
| *krt8* | Q6NWF6 | Keratin, type II cytoskeletal 8 | 0.037 | -2.351 |
| *ctsba* | Q6PH75 | Cathepsin B (Fragment) | 0.002 | -2.362 |
| *calr3b* | A0A0R4IL29 | Calreticulin | < 0.001 | -2.443 |
| *krt18* | Q7ZTS4 | Keratin, type I cytoskeletal 18 | 0.029 | -2.530 |
| *anxa1a* | Q804H2 | Annexin | 0.032 | -2.542 |
| *cnn2* | A0A8M9Q784 | Calponin | < 0.001 | -2.543 |
| *actn1* | A0AB32T375 | Alpha-actinin-1 isoform X6 | 0.007 | -2.647 |
| *atp2a3* | A0A2R8QJH4 | Calcium-transporting ATPase | < 0.001 | -2.648 |
| *htra1b* | A9JRB3 | Serine protease HTRA1B | 0.003 | -2.940 |
| *pfn1* | E7F0A1 | Profilin | 0.011 | -3.122 |
| *ckbb* | Q8AY63 | Creatine kinase | < 0.001 | -3.491 |
| *si:ch211-5k11.8* | Q6ZM17 | Hemoglobin subunit alpha | 0.005 | -3.668 |
| *serpina1* | Q5SPJ4 | Alpha-1-antitrypsin precursor | 0.028 | -4.106 |
| *hbaa1* | Q90487 | Hemoglobin subunit alpha | 0.033 | -4.164 |
| *apoeb* | O42364 | Apolipoprotein Eb | < 0.001 | -4.262 |
| *gapdh* | Q5XJ10 | Glyceraldehyde-3-phosphate dehydrogenase | 0.001 | -4.297 |
| *tfa* | B8JL43 | Serotransferrin | 0.020 | -4.332 |
| *serpina1l* | A0A8N7V0N1 | Serine (Or cysteine) proteinase inhibitor, clade A (Alpha-1 antiproteinase, antitrypsin), member 1, like precursor | 0.018 | -4.504 |
| *hspg2* | A0A8M9PWV8 | Basement membrane-specific heparan sulfate proteoglycan core protein isoform X1 | 0.025 | -4.510 |
| *ba1* | Q90486 | Hemoglobin subunit beta-1 | 0.027 | -4.824 |
| *apoa1b* | A0A0R4IKF0 | Apolipoprotein A-Ib | 0.008 | -5.141 |
| *apoa2* | B3DFP9 | Apolipoprotein A-II | < 0.001 | -5.736 |
| *c3a.1* | A0A8M1P7C1 | Complement C3a.1 precursor | < 0.001 | -5.877 |

**Supplemental Table 9. Significant proteins (p-value < 0.05 and log_2_FC > 1.5 or log_2_FC < -1.5) from *cryaba^-/-^* 4 days post-PBS treated lenses compared to *cryaba^-/-^* 4 days post-H_2_O_2_ treated lenses organized by log_2_FC (n = 73).**

| **Gene group** | **Protein group** | **Protein description** | **p-value** | **Log_2_FC** |
| --- | --- | --- | --- | --- |
| *mdkb* | Q9DDG2 | Midkine | < 0.001 | 3.712 |
| *spon1b* | F1R2Z9 | Spondin-1 | 0.001 | 2.527 |
| *nat16* | E7FET0 | N-acetyltransferase 16 | 0.002 | 1.830 |
| *hspb2* | Q566R8 | Heat shock protein beta-2 | 0.003 | 1.596 |
| *rpl36a* | P61485 | Large ribosomal subunit protein eL42 | 0.002 | 1.592 |
| *uchl3* | Q504C0 | Ubiquitin carboxyl-terminal hydrolase | < 0.001 | 1.571 |
| *gyg1b* | Q6IQP1 | Glycogenin glucosyltransferase | < 0.001 | 1.534 |
| *lctla* | A0AB32TYA7 | Lactase-like a isoform X1 | 0.036 | -1.510 |
| *selenof* | Q802F3 | Selenoprotein F | 0.006 | -1.516 |
| *lim2.3* | Q6DGY5 | Lens intrinsic membrane protein 2.3 | 0.030 | -1.521 |
| *flot1b* | Q6TH07 | Flotillin | 0.008 | -1.526 |
| *vdac3* | A0A8M2BEA1 | Voltage-dependent anion-selective channel protein 3 | 0.017 | -1.528 |
| *psme2* | Q9PUC4 | Proteasome activator complex subunit 2 | 0.014 | -1.534 |
| *etfb* | Q7ZV76 | Electron transfer flavoprotein subunit beta | 0.006 | -1.546 |
| *uqcrb* | Q4VBV1 | Cytochrome b-c1 complex subunit 7 | 0.020 | -1.550 |
| *hspe1* | Q6IQI7 | 10 kDa heat shock protein, mitochondrial isoform 1 | 0.001 | -1.551 |
| *gja3* | Q8JFD2 | Gap junction protein | 0.036 | -1.551 |
| *slc2a1b* | F1R0Q1 | Solute carrier family 2 member 1b | 0.011 | -1.576 |
| *myl6* | F1QWU4 | Myosin light polypeptide 6 isoform X1 | 0.015 | -1.580 |
| *anxa11a* | B8A4V2 | Annexin | 0.034 | -1.593 |
| *psme1* | Q9PTH5 | Proteasome activator complex subunit 1 | 0.001 | -1.600 |
| *enpp6* | Q5BKW7 | Glycerophosphocholine cholinephosphodiesterase ENPP6 | 0.030 | -1.602 |
| *hk1* | Q7ZUM3 | Hexokinase | 0.003 | -1.605 |
| *sfxn3* | A1L257 | Sidoreflexin | 0.006 | -1.623 |
| *lim2.5* | Q5BKW4 | Lens intrinsic membrane protein 2.5 | 0.035 | -1.634 |
| *cox5aa* | Q4VBU7 | Cytochrome c oxidase subunit 5A, mitochondrial | 0.002 | -1.637 |
| *ctsd* | Q8AWD9 | Cathepsin D | 0.011 | -1.640 |
| *ephx1* | Q7ZVB4 | Epoxide hydrolase | 0.010 | -1.660 |
| *atp5f1c* | A0A8M1P526 | ATP synthase subunit gamma | 0.011 | -1.665 |
| *stom* | A0A0R4IGK4 | Stomatin | 0.027 | -1.692 |
| *ankhb* | P58368 | Progressive ankylosis protein homolog B | 0.009 | -1.702 |
| *hspa5* | Q7SZD3 | 78 kDa glucose-regulated protein | 0.003 | -1.703 |
| *dysf* | A0A8M9QH25 | Dysferlin isoform X1 | 0.001 | -1.704 |
| *atp5fa1* | Q08BA1 | ATP synthase subunit alpha | 0.004 | -1.714 |
| *nrcamb* | A0A8M9PV37 | Neuronal cell adhesion molecule | 0.038 | -1.721 |
| *lum* | Q6IQQ7 | Lumican | 0.015 | -1.723 |
| *h2ax* | Q7ZUY3 | Histone H2AX | 0.037 | -1.744 |
| *aldh18a1* | A0A8M3B4R7 | Delta-1-pyrroline-5-carboxylate synthase | 0.012 | -1.758 |
| *tspan33b* | A0A8M2B5Z3 | Tetraspanin | 0.014 | -1.761 |
| *fhl1a* | Q6GQL7 | Fhla protein | 0.045 | -1.768 |
| *gje1b* | A0A8M9P965 | Gap junction epsilon-1 protein-like | 0.009 | -1.796 |
| *atp2b1a* | A0A8M9Q5T9 | Calcium-transporting ATPase | 0.008 | -1.800 |
| *lctlb* | A0A286Y8V0 | Lactase-like b | 0.030 | -1.804 |
| *jam3b* | A3KPA0 | Junctional adhesion molecule 3B | 0.028 | -1.824 |
| *hspd1* | Q803B0 | 60 kDa heat shock protein, mitochondrial | 0.001 | -1.838 |
| *ncstn* | Q5XLR3 | Nicastrin | 0.007 | -1.847 |
| *crygm2d10* | A8E4S8 | Crystallin, gamma M2d10 | 0.004 | -1.856 |
| *hadhb* | Q7ZTH6 | Trifunctional enzyme subunit beta, mitochondrial | 0.004 | -1.876 |
| *hadhab* | A3KMH5 | Enoyl-CoA hydratase | 0.005 | -1.882 |
| *hyou1* | Q7ZUW2 | Hypoxia up-regulated protein 1 | 0.001 | -1.907 |
| *actn1* | A0AB32T375 | Alpha-actinin-1 isoform X6 | 0.004 | -1.917 |
| *pfn1* | E7F0A1 | Profilin | 0.034 | -1.949 |
| *calr3b* | A0A0R4IL29 | Calreticulin | 0.004 | -1.970 |
| *si:dkey-177p2.18* | A0A8M9PTB0 | Phospholipase B1, membrane-associated | 0.015 | -1.988 |
| *cst3* | B8A4C9 | Cystatin C (amyloid angiopathy and cerebral hemorrhage) | 0.020 | -1.988 |
| *npc1* | A0A2R8Q2J0 | NPC intracellular cholesterol transporter 1 isoform X1 | 0.028 | -2.018 |
| *cxadr* | Q90Y50 | Coxsackievirus and adenovirus receptor homolog | 0.010 | -2.119 |
| *hist1h4l* | A3KPR4 | Histone H4 | 0.010 | -2.143 |
| *plod1a* | A8WIN7 | Procollagen-lysine,2-oxoglutarate 5-dioxygenase 1 | < 0.001 | -2.234 |
| *pkp3b* | F1QWE6 | Plakophilin 3b | 0.011 | -2.279 |
| *aldh1a2* | Q90XS8 | Retinal dehydrogenase 2 | 0.017 | -2.280 |
| *fam234a* | A0A8M1PZ20 | Family with sequence similarity 234 member A | 0.020 | -2.306 |
| *anxa1a* | Q804H2 | Annexin | < 0.001 | -2.317 |
| *lgals2a* | Q6IQI2 | Galectin | 0.023 | -2.596 |
| *crygm2d13* | Q6DH12 | Crystallin, gamma M2d13 | < 0.001 | -2.680 |
| *ckbb* | Q8AY63 | Creatine kinase | 0.004 | -2.937 |
| *krt18* | Q7ZTS4 | Keratin, type I cytoskeletal 18 | 0.005 | -3.147 |
| *krt8* | Q6NWF6 | Keratin, type II cytoskeletal 8 | 0.003 | -3.225 |
| *hpx* | Q6PHG2 | Hemopexin | 0.009 | -3.295 |
| *c3a.1* | A0A8M1P7C1 | Complement C3a.1 precursor | 0.009 | -3.567 |
| *crp3* | A0AB32TJM2 | C-reactive protein 3 isoform X1 | 0.020 | -4.001 |
| *apoeb* | O42364 | Apolipoprotein Eb | < 0.001 | -4.429 |
| *serpina1* | Q5SPJ4 | Alpha-1-antitrypsin precursor | 0.005 | -5.100 |

**Supplemental Table 10. Significant proteins (p-value < 0.05 and log_2_FC > 1.5 or log_2_FC < -1.5) from *nrf2^fh318/fh318^* 4 days post-PBS treated lenses compared to *nrf2^fh318/fh318^* 4 days post-H_2_O_2_ treated lenses organized by log_2_FC (n = 11).**

| **Gene group** | **Protein group** | **Protein description** | **p-value** | **Log_2_FC** |
| --- | --- | --- | --- | --- |
| *c1qtnf13* | A0A8M2BHA7 | Uncharacterized protein isoform X1 | 0.004 | 4.053 |
| *col28a1b* | A0A060Q6E4 | Collagen type XXVIII alpha 1 b | 0.002 | 3.882 |
| *col28a1a* | C0H5W1 | Collagen type XXVIII alpha 1 a (Fragment) | 0.001 | 3.662 |
| *nid1a* | F1RAG3 | Nidogen 1a | 0.022 | 3.118 |
| *col18a1a* | A0A8M9QJR2 | Collagen type XVIII alpha 1 chain a isoform X1 | 0.003 | 2.848 |
| *lenep* | A0A8M1P644 | Lens epithelial cell protein LEP503 | 0.015 | 2.391 |
| *tinagl1* | A3KNG7 | Si:dkey-158b13.1 | 0.005 | 2.339 |
| *spon1b* | F1R2Z9 | Spondin-1 | 0.014 | 2.275 |
| *sparcl1* | B3DJ32 | SPARC-like 1 | 0.001 | 2.131 |
| *fthl28* | A8WGK2 | Ferritin | 0.001 | -2.376 |
| *cst14b.2* | A0A2R8RU89 | Cystatin 14b, tandem duplicate 2 | 0.001 | -2.847 |

**Supplemental Table 11. Significant proteins (p-value < 0.05 and log_2_FC > 1.5 or log_2_FC < -1.5) from *cryaba^-/-^;* *nrf2^fh318/fh318^* 4 days post-PBS treated lenses compared to *cryaba^-/-^;* *nrf2^fh318/fh318^* 4 days post-H_2_O_2_ treated lenses organized by log_2_FC (n = 46).**

| **Gene group** | **Protein group** | **Protein description** | **p-value** | **Log_2_FC** |
| --- | --- | --- | --- | --- |
| *nid1a* | F1RAG3 | Nidogen 1a | 0.015 | 3.539 |
| *lenep* | A0A8M1P644 | Lens epithelial cell protein LEP503 | < 0.001 | 3.033 |
| *coch* | Q8AW56 | Cochlin | < 0.001 | 2.777 |
| *tinagl1* | A3KNG7 | Si:dkey-158b13.1 | 0.032 | 2.766 |
| *spon1b* | F1R2Z9 | Spondin-1 | < 0.001 | 2.707 |
| *nedd8* | F1QMF9 | Ubiquitin-like protein NEDD8 | 0.001 | 2.228 |
| *mdkb* | Q9DDG2 | Midkine | < 0.001 | 2.138 |
| *crybgx* | Q45FX8 | BetagammaX-crystallin | < 0.001 | 1.755 |
| *sparcl1* | B3DJ32 | SPARC-like 1 | 0.031 | 1.574 |
| *hspa4l* | X1WBR5 | Heat shock 70 kDa protein 4L | 0.001 | 1.521 |
| *rpl36* | Q6Q415 | Large ribosomal subunit protein eL36 | < 0.001 | 1.517 |
| *tuba8l4* | Q7ZVI6 | Tubulin alpha chain | < 0.001 | -1.518 |
| *palld* | F8W315 | Palladin isoform X1 | 0.001 | -1.534 |
| *zyx* | F1QC03 | Zyxin | 0.002 | -1.540 |
| *hspd1* | Q803B0 | 60 kDa heat shock protein, mitochondrial | < 0.001 | -1.545 |
| *atp2a3* | A0A2R8QJH4 | Calcium-transporting ATPase | 0.001 | -1.547 |
| *rtca* | Q7ZUX5 | RNA 3'-terminal phosphate cyclase | 0.001 | -1.551 |
| *crip1* | Q5BKV7 | Cysteine-rich protein 1 | 0.001 | -1.561 |
| *pcna* | Q9PTP1 | Proliferating cell nuclear antigen | 0.010 | -1.576 |
| *plod2* | A4U7F9 | Procollagen-lysine 5-dioxygenase | 0.004 | -1.611 |
| *hyou1* | Q7ZUW2 | Hypoxia up-regulated protein 1 | < 0.001 | -1.614 |
| *krt8* | Q6NWF6 | Keratin, type II cytoskeletal 8 | 0.001 | -1.627 |
| *fkbp9* | Q6DBZ1 | Peptidylprolyl isomerase | < 0.001 | -1.643 |
| *hsp90b1* | Q7T3L3 | Chaperone protein GP96 | < 0.001 | -1.649 |
| *rrbp1a* | B8A4D7 | Ribosome-binding protein 1a | < 0.001 | -1.684 |
| *jpt1b* | A0A8M2BAY7 | Jupiter microtubule associated homolog 1b isoform X1 | 0.001 | -1.688 |
| *rcc2* | Q6NYE2 | Protein RCC2 homolog | < 0.001 | -1.775 |
| *krt18* | Q7ZTS4 | Keratin, type I cytoskeletal 18 | 0.001 | -1.783 |
| *zgc:162184* | A4QNV6 | Uncharacterized protein LOC559822 | 0.001 | -1.795 |
| *calr3b* | A0A0R4IL29 | Calreticulin | 0.001 | -1.880 |
| *ddx5* | F1Q5L6 | RNA helicase | < 0.001 | -1.884 |
| *hsp70.1* | B0UXR9 | Heat shock cognate 70-kd protein, tandem duplicate 2 | 0.001 | -1.920 |
| *csrp1a* | Q6NZV4 | Cysteine and glycine-rich protein 1 | < 0.001 | -1.932 |
| *cnn2* | A0A8M9Q784 | Calponin | 0.002 | -1.966 |
| *npc1* | A0A2R8Q2J0 | NPC intracellular cholesterol transporter 1 isoform X1 | < 0.001 | -1.970 |
| *efhd2* | Q1LVD7 | EF-hand domain family, member D2 | < 0.001 | -2.005 |
| *hnrnpa0l* | Q7ZU48 | Heterogeneous nuclear ribonucleoprotein A0,-like | < 0.001 | -2.017 |
| *LOC100331639* | A0A8M1RDD4 | Myosin heavy chain, embryonic smooth muscle isoform | < 0.001 | -2.130 |
| *actn1* | A0AB32T375 | Alpha-actinin-1 isoform X6 | < 0.001 | -2.195 |
| *fthl28* | A8WGK2 | Ferritin | < 0.001 | -2.239 |
| *apoeb* | O42364 | Apolipoprotein Eb | 0.004 | -3.044 |
| *ckbb* | Q8AY63 | Creatine kinase | < 0.001 | -3.232 |
| *krt18b* | Q6PHE4 | Keratin, type I cytoskeletal 18b | 0.001 | -3.311 |
| *pfn1* | E7F0A1 | Profilin | < 0.001 | -3.705 |
| *tfa* | B8JL43 | Serotransferrin | < 0.001 | -6.611 |
| *apoa1b* | A0A0R4IKF0 | Apolipoprotein A-Ib | < 0.001 | -6.627 |
