## Supplemental File 8 for "Upregulation of the Unfolded and Mitochondrial Unfolded Protein Responses in Oxidative Stress-Induced Cataract"

**Supplemental Table 12. Significantly enriched GSEA terms from WT 4 days post-PBS treated lenses compared to *cryaba^-/-^* 4 days post-PBS treated lenses, organized by normalized enrichment score (NES).**

| **Go term** | **Description** | **FDR** | **p-value** | **NES** | **Number of proteins with annotation in dataset** |
| --- | --- | --- | --- | --- | --- |
| GO:0015318 | Inorganic molecular entity transmembrane transporter activity | < 1E-03 | < 1E-03 | 2.343 | 30 |
| GO:0007265 | Ras protein signal transduction | 0.006 | < 1E-03 | 2.075 | 44 |
| GO:0044769 | ATPase activity, coupled to transmembrane movement of ions, rotational mechanism | 0.006 | 0.006 | 1.870 | 14 |
| GO:0017111 | Nucleoside-triphosphatase activity | 0.009 | < 1E-03 | 1.832 | 131 |
| GO:0008081 | Phosphoric diester hydrolase activity | 0.009 | < 1E-03 | 1.828 | 10 |
| GO:0050839 | Cell adhesion molecule binding | 0.014 | 0.009 | 1.744 | 10 |
| GO:0003779 | Actin binding | 0.024 | 0.004 | 1.688 | 48 |
| GO:0019900 | Kinase binding | 0.029 | 0.010 | 1.659 | 12 |
| GO:0001664 | G protein-coupled receptor binding | 0.039 | 0.009 | 1.628 | 10 |
| GO:0006952 | Defense response | 0.048 | 0.015 | -1.784 | 16 |
| GO:0016052 | Carbohydrate catabolic process | 0.022 | < 1E-03 | -1.887 | 25 |
| GO:0005975 | Carbohydrate metabolic process | 0.022 | < 1E-03 | -1.891 | 72 |
| GO:0006913 | Nucleocytoplasmic transport | 0.015 | < 1E-03 | -1.935 | 27 |
| GO:0006412 | Translation | 0.012 | < 1E-03 | -1.964 | 133 |
| GO:0050657 | Nucleic acid transport | 0.011 | < 1E-03 | -1.995 | 16 |
| GO:0009116 | Nucleoside metabolic process | 0.005 | < 1E-03 | -2.121 | 22 |
| GO:0034404 | Nucleobase-containing small molecule biosynthetic process | 0.003 | < 1E-03 | -2.149 | 28 |

**Supplemental Table 13. Significantly enriched GSEA terms from *nrf2^fh318/fh318^* 4 days post-PBS treated lenses compared to *nrf2^fh318/fh318^* 4 days post-PBS treated lenses, organized by normalized enrichment score (NES).**

| **Go term** | **Description** | **FDR** | **p-value** | **NES** | **Number of proteins with annotation in dataset** |
| --- | --- | --- | --- | --- | --- |
| GO:0004298 | Threonine-type endopeptidase activity | < 1E-03 | < 1E-03 | -2.108 | 12 |

**Supplemental Table 14. Significantly enriched GSEA terms from *cryaba^-/-^;* *nrf2^fh318/fh318^* 4 days post-PBS treated lenses compared to *cryaba^-/-^;* *nrf2^fh318/fh318^* 4 days post-PBS treated lenses, organized by normalized enrichment score (NES).**

| **Go term** | **Description** | **FDR** | **p-value** | **NES** | **Number of proteins with annotation in dataset** |
| --- | --- | --- | --- | --- | --- |
| GO:0015075 | Ion transmembrane transporter activity | 0.037 | 0.002 | 1.860 | 19 |
| GO:0005996 | Monosaccharide metabolic process | 0.050 | < 1E-03 | -1.866 | 10 |
| GO:0034404 | Nucleobase-containing small molecule biosynthetic process | 0.049 | 0.003 | -1.875 | 12 |
| GO:1901135 | Carbohydrate derivative metabolic process | 0.017 | < 1E-03 | -1.970 | 53 |
| GO:0006399 | tRNA metabolic process | 0.002 | < 1E-03 | -2.111 | 11 |
| GO:1901657 | Glycosyl compound metabolic process | 0.005 | < 1E-03 | -2.117 | 14 |

**Supplemental Table 15. Significantly enriched GSEA terms from WT 4 days post-PBS treated lenses compared to WT 4 days post-H_2_O_2_ treated lenses, organized by normalized enrichment score (NES).**

| **Go term** | **Description** | **FDR** | **p-value** | **NES** | **Number of proteins with annotation in dataset** |
| --- | --- | --- | --- | --- | --- |
| GO:0003735 | Structural constituent of ribosome | < 1E-03 | < 1E-03 | 2.668 | 39 |
| GO:0002183 | Cytoplasmic translational initiation | 0.022 | < 1E-03 | 1.972 | 12 |
| GO:0101005 | Ubiquitinyl hydrolase activity | 0.047 | < 1E-03 | 1.805 | 11 |
| GO:0008081 | Phosphoric diester hydrolase activity | 0.039 | 0.007 | 1.803 | 3 |
| GO:1901137 | Carbohydrate derivative biosynthetic process | 0.046 | < 1E-03 | -1.979 | 28 |

**Supplemental Table 16. Significantly enriched GSEA terms from *cryaba^-/-^* 4 days post-PBS treated lenses compared to *cryaba^-/-^* 4 days post-H_2_O_2_ treated lenses, organized by normalized enrichment score (NES).**

| **Go term** | **Description** | **FDR** | **p-value** | **NES** | **Number of proteins with annotation in dataset** |
| --- | --- | --- | --- | --- | --- |
| GO:0006412 | Translation | < 1E-03 | < 1E-03 | 2.923 | 75 |
| GO:0006413 | Translational initiation | < 1E-03 | < 1E-03 | 2.570 | 23 |
| GO:0002181 | Cytoplasmic translation | < 1E-03 | < 1E-03 | 2.486 | 21 |
| GO:0070647 | Protein modification by small protein conjugation or removal | < 1E-03 | < 1E-03 | 2.344 | 38 |
| GO:0042254 | Ribosome biogenesis | < 1E-03 | < 1E-03 | 2.342 | 16 |
| GO:0008234 | Cysteine-type peptidase activity | < 1E-03 | < 1E-03 | 2.333 | 18 |
| GO:0071826 | Ribonucleoprotein complex subunit organization | 0.001 | < 1E-03 | 2.083 | 23 |
| GO:0043021 | Ribonucleoprotein complex binding | 0.003 | < 1E-03 | 1.996 | 11 |
| GO:0019843 | rRNA binding | 0.004 | < 1E-03 | 1.963 | 5 |
| GO:0016567 | Protein ubiquitination | 0.011 | < 1E-03 | 1.895 | 14 |
| GO:0036094 | Small molecule binding | 0.039 | < 1E-03 | -1.629 | 126 |
| GO:0005509 | Calcium ion binding | 0.036 | 0.007 | -1.644 | 13 |
| GO:0001664 | G protein-coupled receptor binding | 0.033 | 0.009 | -1.655 | 7 |
| GO:0016042 | Lipid catabolic process | 0.039 | 0.007 | -1.747 | 9 |
| GO:0006631 | Fatty acid metabolic process | 0.040 | 0.011 | -1.747 | 9 |
| GO:0022604 | Regulation of cell morphogenesis | 0.038 | 0.002 | -1.755 | 7 |
| GO:0051179 | Localization | 0.029 | < 1E-03 | -1.785 | 132 |
| GO:0048881 | Mechanosensory lateral line system development | 0.025 | 0.002 | -1.799 | 6 |
| GO:0022900 | Electron transport chain | 0.024 | < 1E-03 | -1.818 | 6 |
| GO:0061024 | Membrane organization | 0.023 | 0.004 | -1.823 | 15 |
| GO:0000375 | RNA splicing, via transesterification reactions | 0.022 | < 1E-03 | -1.829 | 20 |
| GO:0007266 | Rho protein signal transduction | 0.018 | 0.002 | -1.866 | 8 |
| GO:0042802 | Identical protein binding | 0.004 | 0.005 | -1.869 | 9 |
| GO:0007265 | Ras protein signal transduction | 0.018 | < 1E-03 | -1.870 | 23 |
| GO:0016050 | Vesicle organization | 0.008 | 0.002 | -1.942 | 17 |
| GO:0006820 | Anion transport | 0.004 | < 1E-03 | -1.999 | 7 |
| GO:0045333 | Cellular respiration | 0.003 | < 1E-03 | -2.061 | 11 |
| GO:0048193 | Golgi vesicle transport | 0.001 | < 1E-03 | -2.124 | 28 |
| GO:0003924 | GTPase activity | < 1E-03 | < 1E-03 | -2.136 | 49 |
| GO:0015318 | Inorganic molecular entity transmembrane transporter activity | < 1E-03 | < 1E-03 | -2.173 | 17 |
| GO:0042060 | Wound healing | < 1E-03 | < 1E-03 | -2.239 | 14 |

**Supplemental Table 17. Significantly enriched GSEA terms from *nrf2^fh318/fh318^* 4 days post-PBS treated lenses compared to *nrf2^fh318/fh318^* 4 days post-H_2_O_2_ treated lenses, organized by normalized enrichment score (NES).**

| **Go term** | **Description** | **FDR** | **p-value** | **NES** | **Number of proteins with annotation in dataset** |
| --- | --- | --- | --- | --- | --- |
| GO:0006412 | Translation | < 1E-03 | < 1E-03 | 2.408 | 75 |
| GO:0042254 | Ribosome biogenesis | 0.001 | < 1E-03 | 2.004 | 19 |
| GO:0002181 | Cytoplasmic translation | < 1E-03 | < 1E-03 | 1.990 | 24 |
| GO:0006413 | Translational initiation | 0.003 | < 1E-03 | 1.905 | 18 |
| GO:0071826 | Ribonucleoprotein complex subunit organization | 0.011 | 0.001 | 1.820 | 24 |
| GO:0009896 | Positive regulation of catabolic process | 0.038 | 0.001 | 1.725 | 11 |
| GO:0006364 | rRNA processing | 0.040 | 0.001 | 1.707 | 7 |

**Supplemental Table 18. Significantly enriched GSEA terms from *cryaba^-/-^;* *nrf2^fh318/fh318^* 4 days post-PBS treated lenses compared to *cryaba^-/-^;* *nrf2^fh318/fh318^* 4 days post-H_2_O_2_ treated lenses, organized by normalized enrichment score (NES).**

| **Go term** | **Description** | **FDR** | **p-value** | **NES** | **Number of proteins with annotation in dataset** |
| --- | --- | --- | --- | --- | --- |
| GO:0043043 | Peptide biosynthetic process | < 1E-03 | < 1E-03 | 2.739 | 135 |
| GO:0002181 | Cytoplasmic translation | < 1E-03 | < 1E-03 | 2.572 | 36 |
| GO:0008234 | Cysteine-type peptidase activity | 0.001 | < 1E-03 | 2.029 | 27 |
| GO:0042255 | Ribosome assembly | 0.003 | < 1E-03 | 1.980 | 16 |
| GO:0042274 | Ribosomal small subunit biogenesis | 0.009 | < 1E-03 | 1.893 | 13 |
| GO:0030218 | Erythrocyte differentiation | 0.018 | 0.004 | 1.821 | 11 |
| GO:0019843 | rRNA binding | 0.041 | 0.007 | 1.703 | 10 |
| GO:0140101 | Catalytic activity, acting on a tRNA | 0.049 | 0.013 | 1.658 | 15 |
| GO:0006820 | Anion transport | 0.022 | < 1E-03 | -1.934 | 15 |
| GO:0031099 | Regeneration | 0.024 | < 1E-03 | -1.954 | 17 |
| GO:0006397 | mRNA processing | 0.017 | < 1E-03 | -2.036 | 39 |
